# Small-molecule-induced species redistribution drives amyloid remodeling along a conserved kinetic landscape

**DOI:** 10.64898/2026.02.22.706957

**Authors:** Radhika V. Nair, Bich Ngoc Tran, Atul N. Parikh, Matthew R. Foreman

**Affiliations:** Institute for Digital Molecular Analytics and Science, Nanyang Technological University, 59 Nanyang Drive, Singapore 636921, Singapore; NTU Institute of Structural Biology (NISB), Nanyang Technological University, Experimental Medicine Building, 59 Nanyang Drive, Singapore 636921; School of Materials Science and Engineering, Nanyang Technological University, 50 Nanyang Avenue, Singapore 639798; School of Electrical and Electronic Engineering, Nanyang Technological University, 50 Nanyang Avenue, Singapore 639798

## Abstract

Proteins in living systems are highly dynamic, continuously populating heterogeneous ensembles of functionally distinct conformational and aggregation states whose relative populations and interconversion rates are governed by an underlying free-energy landscape. Perturbations such as ligand binding, chemical modification, or changes in the environment remodel these assemblies, thereby reshaping protein behavior and function. Understanding how such protein networks are dynamically remodeled is a fundamental challenge in biophysics. Here, we utilize the A*β*_42_ aggregation landscape as a model to demonstrate that remodeling can occur without detectable changes in the apparent activation free-energy barrier, instead arising from system-level redistribution of species along a conserved kinetic pathway. Using EPPS (4-(2-Hydroxyethyl)-1-piperazinepropanesulfonic acid) as a model modulator, temperature-dependent kinetic analysis reveals an invariant apparent activation free energy, suggesting that the underlying energetic framework remains unchanged even as the system is reshaped. We find that EPPS modulates A*β*_42_ disaggregation in a concentration-dependent manner. At intermediate concentrations, enhanced redistribution toward soluble species is observed, whereas at higher concentrations, modulator self-association into supramolecular clusters is associated with reduced net disaggregation and partial recovery of aggregation signatures. Thermal disruption of these assemblies restores activity, consistent with a role of modulator self-association in governing the observed behavior. Furthermore, our experiments indicate that this disaggregation process is reduced in crowded environments compared to dilute conditions. Together, these results support a biophysical framework in which amyloid remodeling is driven by species redistribution along a conserved kinetic landscape, where the extent of disaggregation is determined by modulator concentration-dependent partitioning of protein species, modulator self-association, and environmental constraints.

**Significance:** An emerging view of protein function recognizes that proteins exist as dynamic networks of interconverting conformational and aggregation states. Binding interactions and environmental perturbations remodel these networks, thereby altering protein behavior. Amyloid aggregates associated with Alzheimer’s disease exemplify such dynamic systems, yet how small molecules remodel them remains poorly understood. Using A*β*_42_ as a model, we show that small molecules remodel amyloid networks by redistributing protein populations in a concentration-dependent manner. This redistribution is modulated by self-association of the small molecule, molecular crowding, and temperature. These findings establish population redistribution within dynamic amyloid networks as a mechanism for remodeling protein assemblies, offering new insights into controlling pathological aggregation.

## Introduction

Amyloid-*β* (A*β*) peptides, 39-43 amino acids in length (~4 kDa), are generated by sequential cleavage of the amyloid precursor protein (APP, ~ 120 kDa) by *β*- and *γ*-secretases. ^1^ Owing to the poor solubility in aqueous media, they readily self-associate both *in vivo* and *in vitro* producing structurally heterogeneous mixtures of higher-order, mesoscopic aggregates. ^2,3^ These aggregates comprise a diverse variety of species, including *β*-sheet-rich oligomers, spherical non-*β*-sheet aggregates, micelles, protofibrils, and mature fibrils. Collectively, these species span a broad conformational phase space: some correspond to distinct on-pathway intermediates en route to fibril formation, whereas others, such as non-*β*-sheet aggregates that transiently sequester monomers or oligomers, represent off-pathway intermediates residing in local free-energy minima along side branches of the amyloid aggregation landscape. ^4,5^

Rather than existing as isolated entities, these structurally diverse aggregates are dynamically interconnected, collectively forming an interconnected reaction network that underlies amyloid polymorphism (Figure 1). Each A*β* aggregate species represent local nodes within an elaborate network linking monomers to mature fibrils across a rugged free-energy landscape. This network consists of multiple competing pathways, with nodes corresponding to local free-energy minima associated with distinct aggregation products. Within this reaction network, fibril formation proceeds predominantly along a canonical nucleation-dependent polymerization pathway. This self-assembly process comprises two phases: (1) a lag phase, during which stochastic association of soluble monomers overcomes an energy barrier to form a stable critical nucleus, and (2) an elongation phase, in which monomer (or oligomer) addition to fibril seeds is thermodynamically favored (Δ*G <* 0). ^6^ Beyond this dominant route, secondary nucleation and oligomer-fibril interactions introduce alternative side pathways that further diversify the aggregation free-energy landscape. ^7–9^

**Figure 1.**
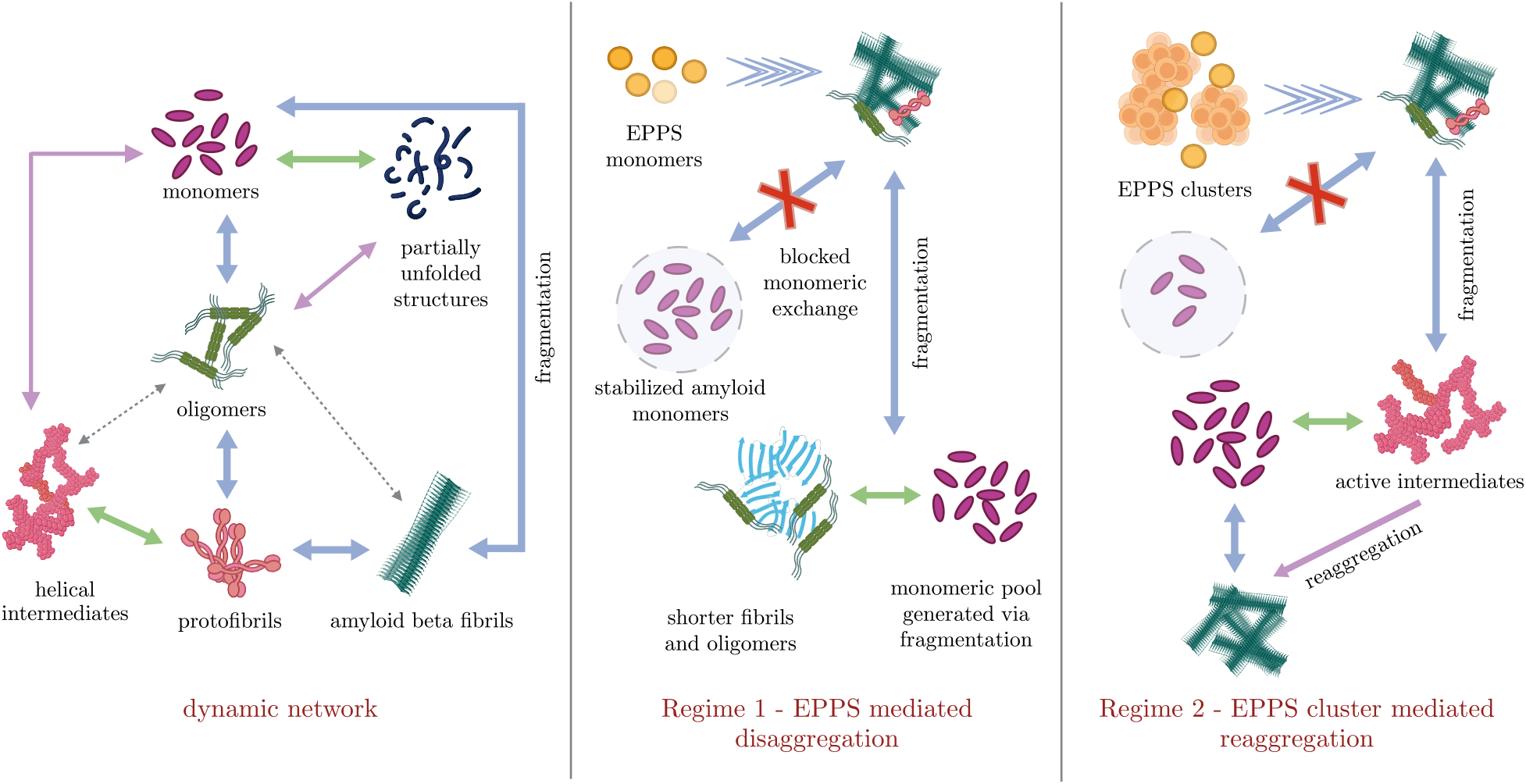
EPPS-mediated modulation of amyloid assembly. Initially in the absence of EPPS, fully grown amyloid fibrils exist in dynamic equilibrium with a network of aggregates and monomers (left). At low concentrations of EPPS (Regime 1), EPPS molecules stabilize amyloid monomers, preventing monomeric exchange which promotes fragmentation of fibrils into shorter fibrils or oligomers generating a new pool of amyloid monomers (middle). At higher EPPS concentrations, EPPS molecules self cluster, reducing the availability of free EPPS monomers to stabilize amyloid monomers. This results in slight fragmentation of amyloid fibrils producing active and unstable intermediates that readily reaggregate due to low free energy barriers (right).

In living systems, the steady-state distribution of higher-order amyloids under any condition is shaped not only by aggregation but also by opposing disaggregation processes. A*β* aggregates are continually counterbalanced by these competing pathways, which are often mediated by molecular chaperons such as HSP70 and clusterin. Through fibril fragmentation and/or monomer release, they generate additional mesoscopic aggregates. A dynamic balance between the two - aggregation and disaggregation - thus governs amyloid turnover. ^10^ Disruption of this finely tuned balance, such as through proteostatic failure, oxidative stress, or aberrant membrane interactions, can shift the equilibrium toward the accumulation of neurotoxic species. ^11^ Conversely, deliberate exogenous perturbations that modulate aggregation or disaggregation kinetics can remodel the reaction network, redistributing the populations of mesoscopic amyloid aggregates and potentially restoring proteostasis. We therefore hypothesize that therapeutic strategies that modulate population distributions within this dynamic network, rather than focusing on individual species alone, may offer improved control over protein self-assembly.

Viewed through this dynamic network perspective, existing therapeutic approaches can be broadly classified into two strategies: early suppression of aggregation processes ^12^ and enhancement of disaggregation pathways. ^13^ For the former, small molecules such as polyphenols (e.g., EGCG and curcumin), ^14–16^ Congo red, ^17^ benzofurans, ^18^ and stilbenes ^19^ have proved promising because they interfere with monomer-monomer interactions. Other approaches that inhibit aggregation employ molecular chaperones, ^20–22^ metal chelators, ^23^ *β*-sheet-breaking peptides, ^24,25^ or monoclonal antibodies. ^26^The latter approach, which targets preformed aggregates to restore proteostasis through active disassembly, ^13^ includes several natural products and engineered systems. Representative examples include natural products such as flavonols, which remodel amyloids via hydroxyl-mediated disruption of *β*-sheet stacking ^27^ and engineered systems such as polymeric micelles, which generate reactive oxygen species and promote oxidative fibril cleavage, ^28^ molecular chaperones (heat shock proteins), which drive disassembly through ATP-dependent mechanisms, ^29^ proteolytic fragmentation, ^30^ or isomerase-mediated remodeling, ^31^ and humanized antibodies such as Crenezumab ^32^ and Gantenerumab, ^33^ which selectively clear aggregated species.

In addition, sequestration or stabilization of monomeric amyloid species has emerged as an alternative strategy for suppressing aggregation by reducing the population of aggregation-competent molecules. ^34–36^ Notably, the small molecule EPPS (4-(2-hydroxyethyl)-1-piperazinepropanesulfonic acid) has been reported both as a protector of monomeric A*β* species and as an effective disaggregating agent for mature amyloid fibrils through a range of proposed interactions. ^37–39^ Amyloid systems, however, exist as heterogeneous ensembles spanning monomers, oligomers, protofibrils, and fibrils, raising the question of how EPPS, through its interactions with multiple amyloid species, perturbs the aggregation landscape of such a dynamic network. Here, we investigate how EPPS, a putative small molecule disaggregase with a molecular weight of 252 Da, reshapes the aggregation landscape of higher order amyloid assemblies. Contrary to the prevailing view that EPPS acts solely through direct and local interactions with amyloid fibrils via its amino sulfonic acid moiety, ^37,40,41^ our results suggest that EPPS acts as a system-level modulator by redistributing the populations of amyloid species. This redistribution is strongly concentration dependent. At low concentrations, EPPS promotes disaggregation, whereas at higher concentrations it favors the formation of distinct higher-order aggregates. This biphasic behavior appears to arise from the concentration-dependent physical state of EPPS itself. At low concentrations, EPPS exists predominantly as monomers and small clusters, whereas at higher concentrations it self-associates into larger mesoscopic assemblies, consistent with supramolecular organization reported for polyphenols and other aromatic small molecules. ^42–44^ Temperature-dependent kinetic measurements further show that aggregate disruption occurs only within the low-concentration disaggregation regime. Because the resulting products neither recover *β*-sheet structure nor regain aggregation activity, we propose that EPPS promotes the sequestration and stabilization of lower-order amyloid species, thereby limiting their re-entry into aggregation pathways (Figure 1). Consistent with this model, the apparent free energy remains largely unchanged during remodeling and retains values close to the intrinsic transition free energy. ^45^ This energetic invariance indicates that EPPS does not fundamentally alter the underlying reaction barrier but instead redistributes amyloid populations into kinetically stable, trapped states. More broadly, our findings suggest that the efficacy of small molecule modulators depends not only on their molecular structure and how they interact with mature fibrils but also on their own mesoscopic organization. In addition, they reveal an underappreciated role of molecular crowding in shaping ensemble-level amyloid-drug interactions.

## Results and Discussion

### Cooperativity in EPPS-Mediated Amyloid Disassembly

We began by preparing mature A*β*_42_ fibrils, establishing a controlled reference state essential for interpreting perturbations. The fibrils were aged until they reached a steady-state assembly, confirmed by a plateau in Thioflavin T (ThT) fluorescence (Supplementary Figure S1). Dynamic light scattering (DLS) (Supplementary Figure S1) and transmission electron microscopy (TEM) (Figure 3(e)) further indicated a predominantly intact fibrillar population with no significant changes in size distribution or morphology under the measurement conditions. Subsequently, we assessed how EPPS interacts with A*β* fibrils, producing measurable changes in aggregation-disaggregation behavior.

Alterations in the cross-*β* structure are expected to be reflected in ThT fluorescence intensity, ^46^ allowing us to monitor structural changes in real time. ThT kinetics revealed two distinct regimes of EPPS action. At low EPPS concentrations (2–18 *µ*M; Regime 1), fluorescence decreased monotonically, indicating concentration-dependent disassembly of cross-*β* structures (Figure 2(a)). Fitting the fluorescence intensity traces I(t) with a Hill model, 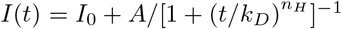, yielded a half-time (*k*_*D*_) that decreased, and a Hill coefficient (*n*_*H*_) that initially increased with EPPS concentration (Figure 2(b)). These trends indicate an increasing sensitivity of the disassembly response to EPPS with increasing concentration, accompanied by an initially enhanced cooperativity of the observed response. As EPPS approaches ~ 16 *µ*M, the cooperativity declines, marking a transition in the system’s response regime. Hill analysis of the steady-state data (from time-based ThT measurements for Regime 1, Supplementary Figure S2 and the binding isotherm (Supplementary Figure S3) further support this transition, showing near-complete saturation of the observed response close to the transition regime.

**Figure 2.**
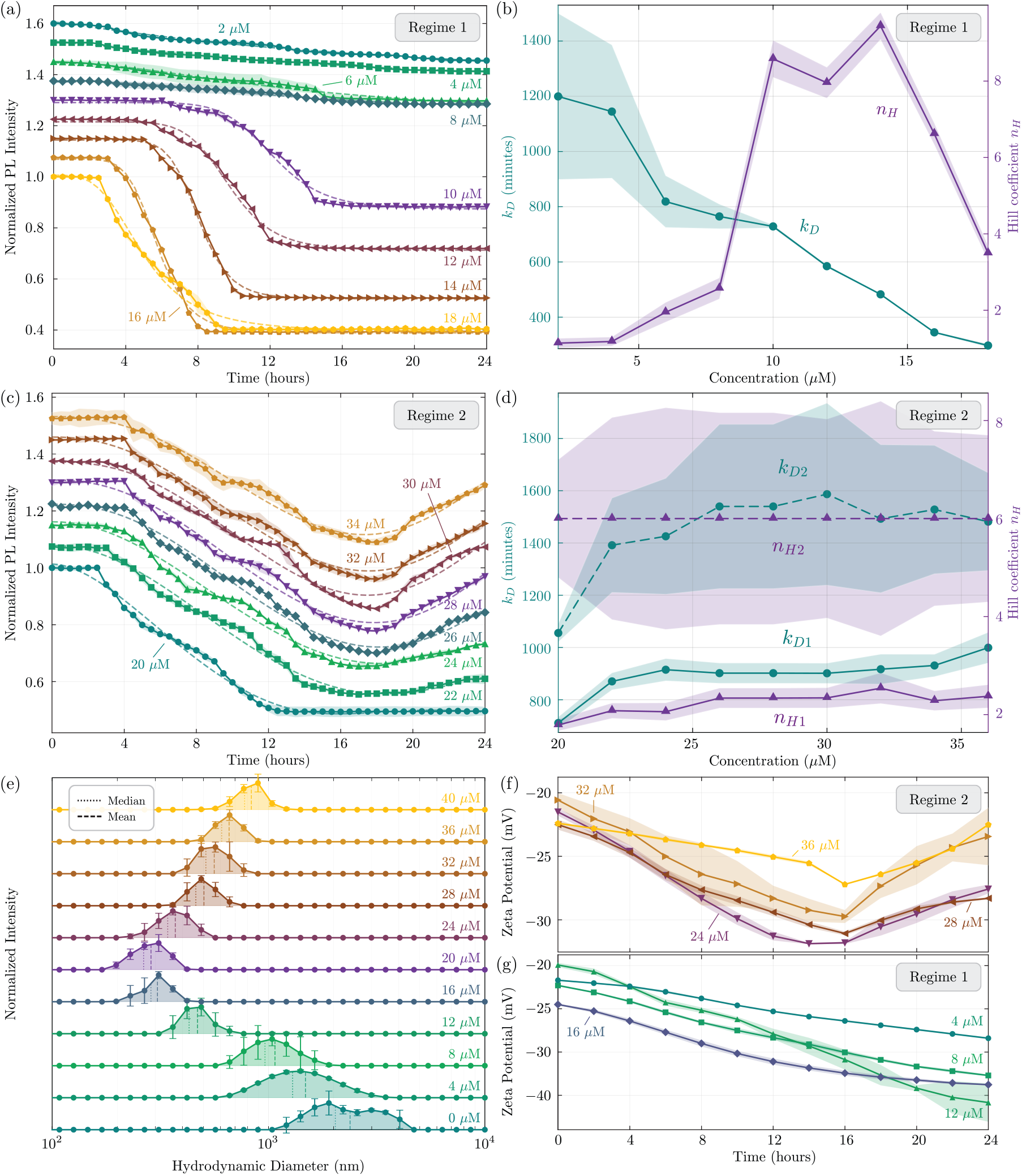
Kinetics and physicochemical characterization of EPPS treated A*β*. (a) observed ThT fluorescence intensity as a function of time for lower concentrations of EPPS (2 *µ*M to 18 *µ*M - Regime 1) and (b) corresponding extracted kinetic parameters *k*_*D*_ and *n*_*H*_ extracted using Hill fit. Similarly, (c) ThT fluorescence intensity profiles and (d) kinetic parameters *k*_*D*1_, *n*_*H*1_, *k*_*D*2_, *n*_*H*2_ extracted using a double Hill fit for higher EPPS concentrations (20-40 *µ*M, Regime 2). Note, each trace has been vertically displaced in and (c) by 0.075 for clarity. (e) DLS measurements showing changes in amyloid size upon treatment with various concentrations of EPPS; dashed lines indicate the mean, while dotted lines represent the median from three measurements. (g) and (f) show surface charge variations of A*β* during EPPS treatment studied using zeta potential measurements for Regime 1 and Regime 2 respectively. Error bars in (e) and shaded bands (others) represent the standard deviation from three replicate measurements.

**Figure 3.**
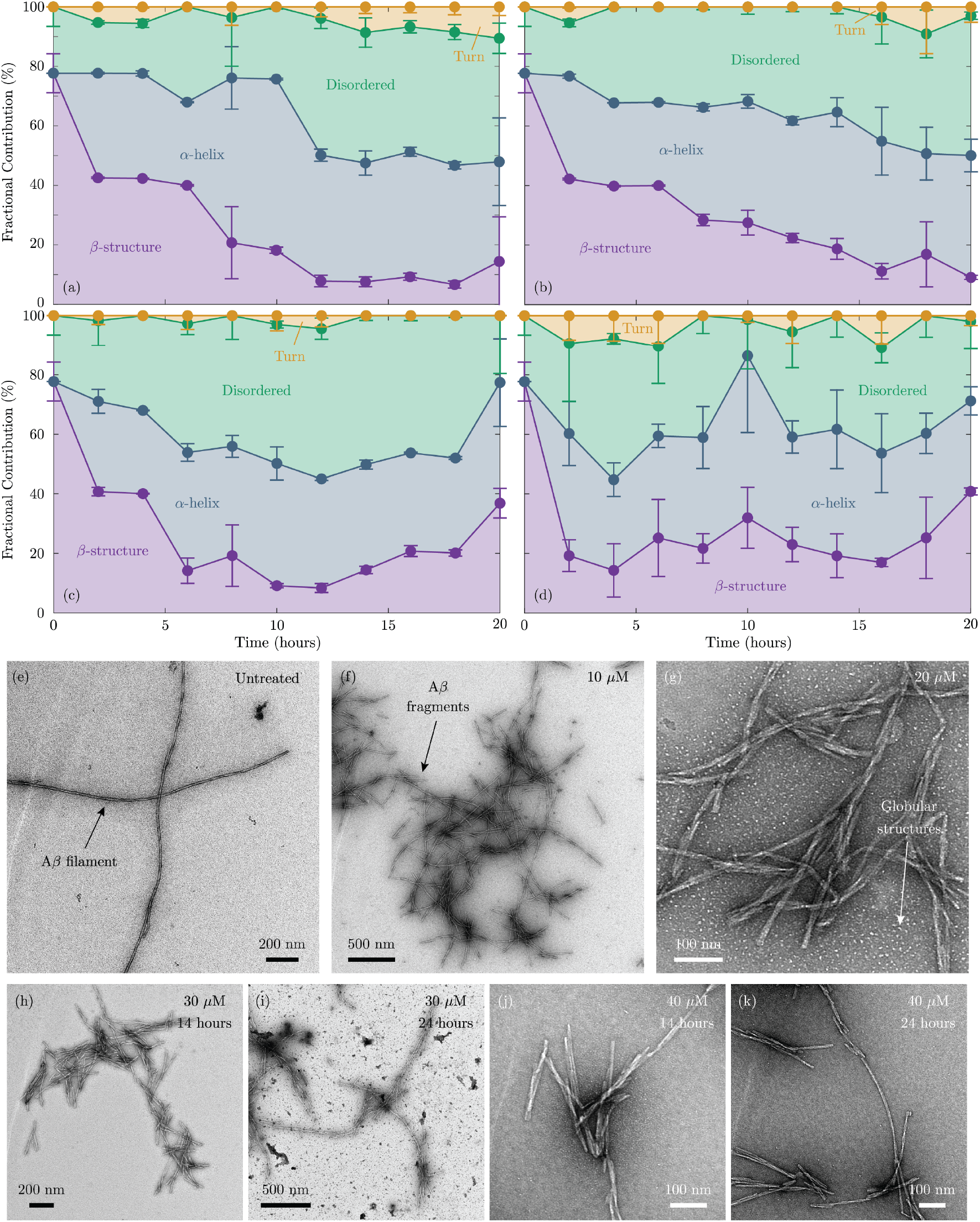
Structural analysis of EPPS treated A*β*. (a)-(d): FTIR analysis performed as a function of time for various EPPS concentrations showing structural transitions during disaggregation and reaggregation processes. Stacked plots show fractional contribution of different amyloid secondary structures over time upon treatment with EPPS at (a) 10 *µ*M, (b) 20 *µ*M, (c) 30 *µ*M, and (d) 40 *µ*M. Error bars indicate standard deviation from 3 independent replicates. (e)-(k) : TEM images of (e) untreated A*β* fibril samples, and samples treated with EPPS at (f) 10 *µ*M, (g) 20 *µ*M, (h) 30 *µ*M at 14 h, and (i) 24 h, and 40 *µ*M at (j) 14 h and (k) 24 h.

At higher concentrations (20–40 *µ*M; Regime 2), the kinetics became biphasic (Figure 2(c)), with an initial decrease in ThT signal followed by a partial recovery. These dynamics were described using a double-Hill model of the form

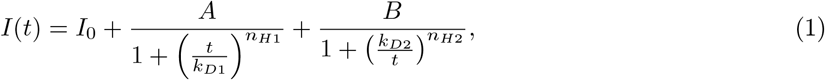

where *I*(*t*) denotes the measured intensity at time *t, I*_0_ is the initial intensity, *k*_*D*1_ and *k*_*D*2_ are characteristic time constants, and *n*_*H*1_ and *n*_*H*2_ are the corresponding Hill coefficients describing the steepness of the observed transition. The two time-dependent terms capture concurrent contributions to the overall ThT response, corresponding to an initial decrease and a subsequent partial recovery of signal.

The fitted parameters (Figure 2(d)) indicate that both phases exhibited cooperative features, with the second phase showing higher apparent cooperativity (*n*_*H*2_ *> n*_*H*1_ *>* 1). The slower timescale of the second process (*k*_*D*2_ ≫ *k*_*D*1_) and the weak dependence of the fitted parameters on EPPS concentration suggest that the observed biphasic behavior is not governed by EPPS concentration alone but reflects intrinsic system kinetics under these conditions. Together, these results indicate a transition in the kinetic response from a predominantly single-phase regime at low concentrations to a biphasic regime at higher concentrations.

Dynamic light scattering (DLS) measurements provided structural context for the kinetic transitions inferred from the double-Hill analysis. After 24 h of incubation, untreated A*β*_42_ fibrils exhibited a large hydrodynamic size of ~ 2.4 *µ*m (Figure 2(e)). The average hydrodynamic diameter of the *Aβ*_42_ assemblies decreased over the EPPS concentration range of 0-20 *µ*M, a trend consistent with the concentration-dependent reduction in ThT signal observed in Regime 1. The mean hydrodynamic diameter (*D*_*H*_) decreased from ~ 2.5 *µ*m to ~ 250 nm, indicating a shift toward smaller scattering populations under these conditions. At higher EPPS concentrations (20–40 *µ*M; Regime 2), the size distribution shifted toward larger values, with *D*_*H*_ increasing to ~ 800 nm. This trend was consistent with the biphasic kinetics observed in the ThT measurements, where an initial decrease in signal was followed by a partial recovery. Overall, the changes in hydrodynamic size distributions reflect a concentration-dependent redistribution of scattering populations across the EPPS concentration range, consistent with the existence of two distinct kinetic regimes.

Electrostatic measurements further substantiated these kinetic distinctions (Figures 2(f) and 2(g)). Zeta potential analysis across EPPS concentrations ranging from 6 to 18 *µ*M revealed a systematic shift toward more negative surface potentials (Figure 2(g)). This trend indicates a concentration-dependent alteration in the surface electrostatic properties of the fibrillar assemblies. The increasing negative surface potential with EPPS concentration is consistent with the disassembly behavior observed in Regime 1, suggesting a redistribution of surface-exposed charged species as fibrillar structures are progressively perturbed.

Beyond 18 *µ*M EPPS, the zeta potential exhibited a biphasic response that paralleled the behavior observed in the double-Hill kinetics (Figure 2(f)). Initially, the surface potential became more negative, coinciding with the rapid decrease in ThT signal associated with the first kinetic phase (*k*_*D*1_). Over extended incubation, the potential gradually shifted toward less negative values, corresponding to the slower second phase (*k*_*D*2_). Together with the ThT and DLS measurements, these electrostatic changes indicate that the two kinetic regimes are accompanied by distinct changes in the physicochemical properties of the A*β* assemblies. Low EPPS concentrations were associated with decreasing ThT intensity, reduced hydrodynamic size, and increasingly negative surface potentials, whereas higher EPPS concentrations produced biphasic kinetic behavior accompanied by partial recovery of both aggregate size and surface electrostatic properties over time. These observations collectively support the existence of two concentration-dependent regimes of EPPS activity that differ in their kinetic, structural, and electrostatic characteristics.

To gain insight into the structural reorganization of amyloid species induced by EPPS, we employed FTIR spectroscopy. FTIR analysis of EPPS-treated amyloid revealed concentration-dependent remodeling of the A*β* secondary-structure distribution, providing direct evidence of changes in the cross-*β* architecture. Fully matured A*β*_42_ fibrils established a structural reference, with the Amide I region (1600–1700 cm^−1^) deconvolved using pseudo-Voigt fitting optimized by AIC/BIC analysis. ^47^ We resolved two principal components: a dominant *β*-sheet band (1610–1635 cm^−1^) and a minor contribution from disordered or monomeric segments (1635–1650 cm^−1^). Quantitative peak-area analysis indicated a *β*-sheet content of ~ 80%, defining a baseline structural state for subsequent comparison (Supplementary Figure S4(a)).

Time-resolved FTIR analysis of A*β*_42_ samples treated with 10–40 *µ*M EPPS captured distinct, concentration-dependent changes in secondary structure (Figures 3(a) to 3(d)). A representative spectrum from the time-resolved FTIR measurements is shown in Supplementary Figure S4(b). In Regime 1 (10-20 *µ*M), the spectra exhibited a progressive loss of *β*-sheet signal within the first 2 h, decreasing from ~ 78% to ~ 40%, concurrent with the emergence of a pronounced *α*-helical component (Figures 3(a) and 3(b)). Over extended incubation times (up to 16 h), the *β*-sheet content further declined to ~ 10%, while the combined *α*-helical and disordered fractions increased to ~ 85%. These observations indicate that, at low EPPS concentrations, the reduction in *β*-sheet structure is accompanied by a substantial redistribution toward non-*β* conformations. The resulting populations may comprise a mixture of monomeric, oligomeric, and structurally disordered species, although the present measurements do not resolve their precise molecular identities.

At higher EPPS concentrations, the structural evolution became more complex, mirroring the biphasic behavior observed in Regime 2. Samples treated with 30 *µ*M EPPS showed an initial decrease in *β*-sheet content to ~ 10%, followed by a partial recovery to ~ 35% at longer times (Figure 3(c)). The combined non-*β* population reached a maximum of ~ 90% before decreasing to ~ 65%, indicating a time-dependent redistribution of secondary-structure populations. At 40 *µ*M, the structural profiles exhibited heterogeneous, non-monotonic fluctuations in both *β*-sheet and disordered fractions, reflecting increased temporal variability in the structural response. Overall, the FTIR measurements reveal a concentration-dependent shift in structural behavior, with lower EPPS concentrations associated with extensive loss of *β*-sheet content and higher concentrations exhibiting partial recovery and greater fluctuations in secondary-structure populations over time.

These FTIR observations are broadly consistent with the kinetic and physicochemical measurements. During the first 3–4 h, FTIR and zeta potential detect early changes in secondary structure and surface electrostatics, including a reduction in *β*-sheet content and increasingly negative surface potentials, prior to substantial changes in ThT fluorescence. This temporal offset suggests that structural and electrostatic rearrangements precede the larger cross beta changes detected by ThT. The subsequent partial recovery of *β*-sheet content at intermediate concentrations coincides with increases in hydrodynamic size measured by DLS and with the second kinetic phase identified by the double-Hill analysis, while the heterogeneous behavior observed at higher EPPS concentrations is reflected across all measurement modalities. Taken together, these observations support a concentration-dependent redistribution of A*β* structural populations, accompanied by corresponding changes in aggregate size and surface electrostatic properties.

Finally, negative-stain transmission electron microscopy (TEM) provided direct visualization of the morphological changes associated with the kinetic and spectroscopic observations. Untreated A*β*_42_ fibrils appeared as long, unbranched filaments with lengths of ~ 1.5–4.5 *µ*m (Figure 3(e)), consistent with mature fibrillar assemblies. Following EPPS treatment, TEM revealed a concentration-dependent fragmentation pattern that closely mirrored the behavior observed in Regime 1. At 10 *µ*M EPPS (Figure 3(f)), fibril lengths were substantially reduced to ~ 600–1000 nm, while at 20 *µ*M EPPS (Figure 3(g)), most filaments shortened further to ~ 200–500 nm. In addition to these shortened fragments, numerous globular species smaller than ~ 10 nm were observed, indicating the presence of smaller assemblies generated during EPPS treatment. This concentration (20 *µ*M) corresponds to the point of maximal disaggregation efficiency identified by ThT, DLS, and zeta potential measurements. Morphological evolution at higher EPPS concentrations reflected the more complex behavior observed in Regime 2. Samples treated with 30 *µ*M EPPS (Figure 3(h)) and 40 *µ*M EPPS (Figure 3(j)) initially displayed pronounced fragmentation after 14 h, with typical lengths of ~ 250–500 nm. Prolonged incubation to 24 h led to the appearance of extended fibrillar structures (Figure 3(i) and Figure 3(k)), ranging from ~ 800 nm to 2 *µ*m, indicating a partial recovery of elongated fibrillar morphologies. This morphological evolution is consistent with the second kinetic phase identified by the double-Hill analysis. The observed trend is further supported by DLS measurements after 24 h incubation, which show an increase in D_*H*_ from ~ 600 nm at 30 *µ*M to ~ 900 nm at 40 *µ*M, together with the partial recovery of *β*-sheet content detected by FTIR.

Longer incubations (4–6 days) produced no further detectable changes in fibril length or morphology relative to the 24 h endpoint, indicating that the EPPS-induced structural remodeling had largely stabilized over the timescale examined (Supplementary Figure S5). This morphological persistence is consistent with the plateau observed in the ThT and DLS measurements, suggesting that the major structural and size redistribution processes occur within the first 24 h under the conditions studied.

The incomplete disaggregation and plateauing behavior observed at higher EPPS concentrations could not be readily explained from the fibril structural data alone, as both Regime 1 and Regime 2 ultimately produced substantial reductions in *β*-sheet content. This observation suggested that factors other than the final amyloid structure might influence EPPS activity at elevated concentrations. Given the zwitterionic nature of EPPS and the concentration-dependent changes in disaggregation behavior, we hypothesized that EPPS may undergo self-association in solution, thereby altering the availability of free EPPS molecules. To investigate this possibility, we next examined the concentration-dependent solution behavior of EPPS using DLS.

### EPPS Self-Association

DLS measurements provided insight into the concentration-dependent solution behavior of EPPS (Figure 4). At concentrations below 20 *µ*M, EPPS exhibited predominantly small hydrodynamic species with radii below 100 nm. Above this range, a marked increase in hydrodynamic size was observed, reaching ~ 400–500 nm at 30–40 *µ*M, indicating a shift toward larger scattering populations.

**Figure 4.**
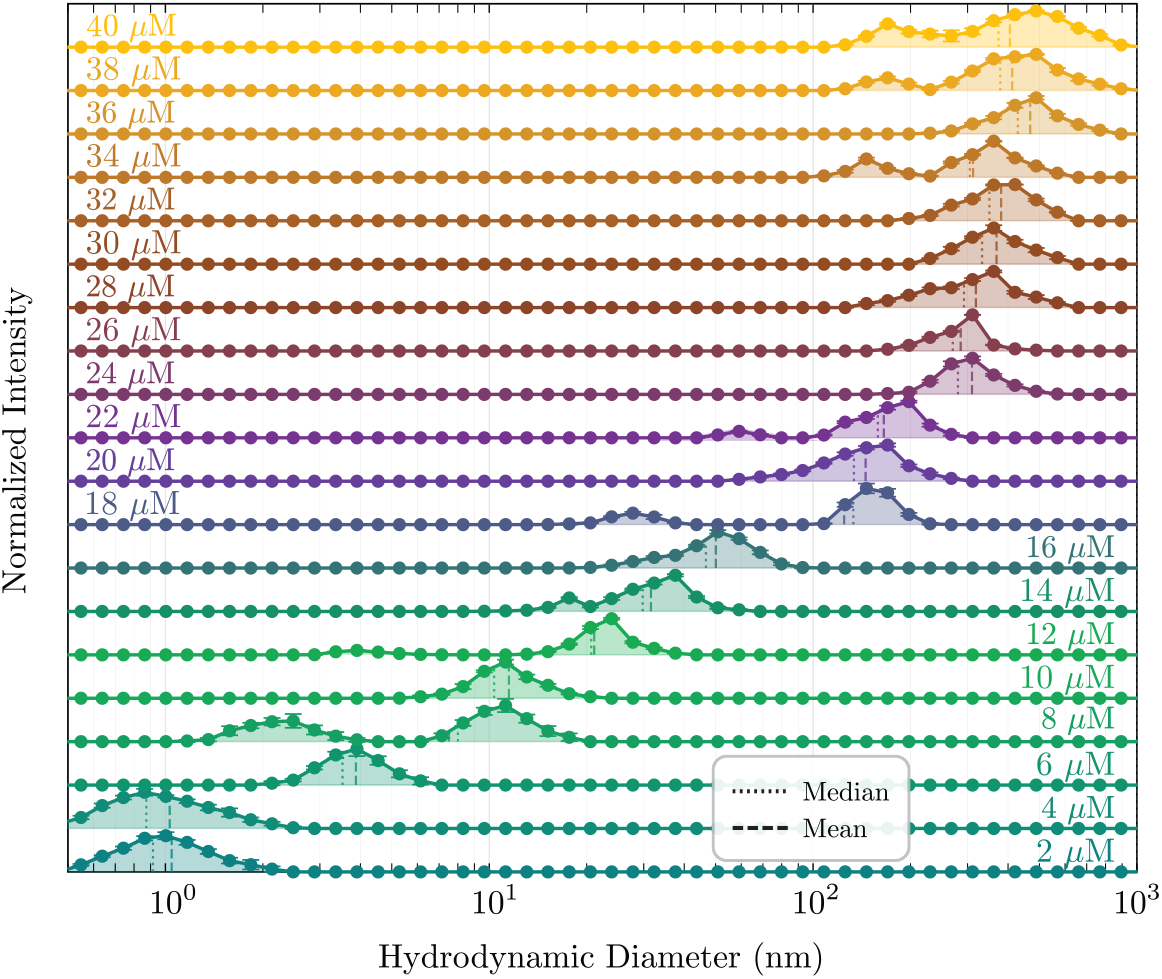
EPPS clustering dynamics as a function of concentration. DLS measurements of EPPS clustering after 24 h incubation across a range of concentrations from 2 to 40 *µ*M. At low EPPS concentrations (2-10 *µ*M), the clusters remain very small, with average sizes below 10 nm, close to the monomeric state. As the concentration increases beyond 10 *µ*M, larger clusters begin to form, reaching sizes of 600-800 nm at the highest EPPS concentrations. The corresponding DLS spectra are presented, with each trace vertically offset for clarity. Dashed and dotted lines indicate the mean and median cluster sizes, respectively, and error bars represent the standard deviation from three independent replicates.

To further examine the distribution of species, DLS intensity profiles were analyzed using a scattering-based framework to estimate relative contributions of different size populations (Supplementary Section 6). This analysis suggests a concentration-dependent redistribution from smaller to larger EPPS-associated species, with the crossover occurring in the same concentration range that separates Regime 1 and Regime 2 in the amyloid experiments. Under high-salt conditions (200 mM NaCl), larger apparent particle sizes were observed across all concentrations (Supplementary Figure S6), indicating that ionic strength influences the extent of EPPS-associated clustering.

Similar behavior was observed for the zwitterionic compound HEPES, which is structurally analogous to EPPS and shares comparable charge-balanced (zwitterionic) character, although it differs in the length of its hydrocarbon chain. Specifically, HEPES exhibited a comparable two-regime response, albeit with a shifted transition threshold (Supplementary Figure S7(a) to S7(d)). DLS measurements (Supplementary Figure S7(e)) further revealed concentration-dependent clustering of HEPES, analogous to that observed for EPPS. Collectively, these observations suggest that concentration-dependent self-association is an important contributor to the concentration-dependent behavior of these small-molecule modulators in amyloid systems. Under the conditions examined, the greatest reduction in aggregation-associated signatures was observed at intermediate concentrations, whereas higher concentrations were associated with diminished net disaggregation and partial recovery of aggregation-associated signatures over time.

To determine whether the concentration-dependent effects of EPPS on A*β*_42_ aggregation landscapes are governed by the existence of supramolecular EPPS clusters, we performed DLS measurements (Figure 5(a)) at elevated temperatures and further thermodynamic studies. We hypothesized that if the inhibitory potency of EPPS is determined by the balance between its self-associated (clustered) and monomeric states, increasing the system’s thermal energy would shift this equilibrium, thereby affecting the availability of species potentially involved in fibril disassembly or the formation of active intermediates. We performed DLS measurements across four concentrations (10 *µ*M and 20 *µ*M in regime 1; 30 *µ*M and 40 *µ*M in regime 2) of EPPS over a temperature range of 30°C to 45°C, in 5°C increments. At all concentrations, the mean hydrodynamic radius of the EPPS species exhibited a monotonic decrease with increasing temperature, indicating a thermally driven dissociation of the supramolecular clusters. For the 10–30 *µ*M samples, the species transitioned to a fully monomeric state at 45 °C, with the mean hydrodynamic diameter ranging from 1.52 ± 0.35 nm to 2.18 ± 0.10 nm. In contrast, at 40 *µ*M, a population of small clusters with an average size of 11.48 ± 0.35 nm persisted even at 45°C. These data indicate that the thermal response of EPPS clusters is concentration-dependent. While elevated temperatures reduced the hydrodynamic size of the assemblies, clustered species remained detectable at higher EPPS concentrations, suggesting that the observed cluster population is governed by the interplay between concentration-dependent association and thermal dissociation. Full DLS spectra are provided in Supplementary Figure S8.

**Figure 5.**
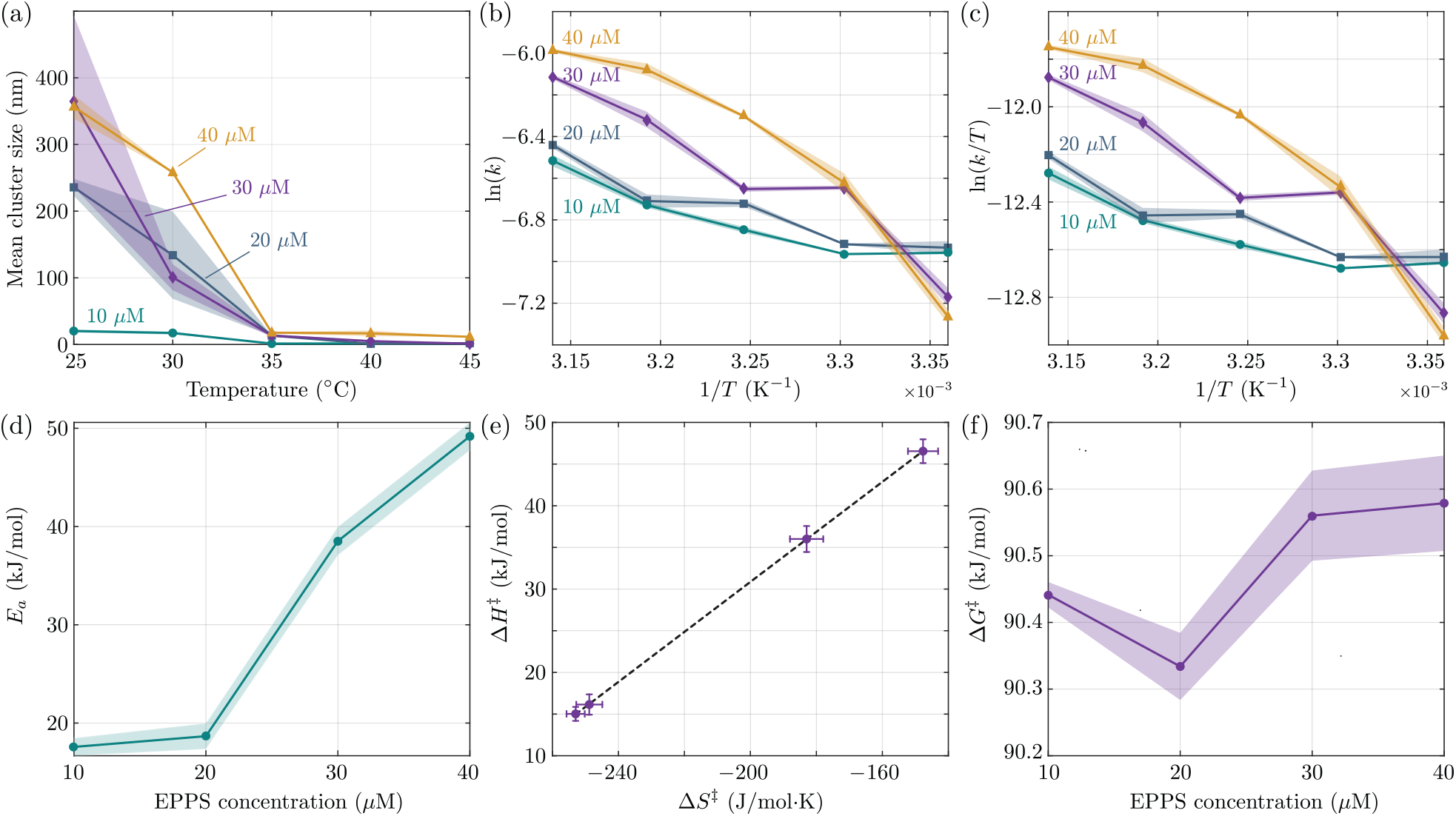
Temperature-dependent EPPS self-association and thermodynamic characterization of EPPS-induced *Aβ*_42_ disaggregation. (a) DLS-measured mean hydrodynamic diameter of EPPS species at 10, 20, 30, and 40 *µ*M across temperatures from 25°C to 45°C. (b) Arrhenius plots used for the calculation of apparent activation energy (*E*_*a*_). (c) Eyring-Polanyi plots used for the calculation of apparent thermodynamic parameters, including activation enthalpy (Δ*H*^*‡*^), activation entropy (Δ*S*^*‡*^), and Gibbs free energy (Δ*G*^*‡*^). Shaded regions represent the uncertainty in the measurements derived from bootstrap analysis. (d) Apparent activation energy (*E*_*a*_) derived from Arrhenius plots. (e) Enthalpy-entropy compensation plot (Δ*H*^*‡*^ vs. Δ*S*^*‡*^) calculated from Eyring-Polanyi plots. (f) Gibbs free energy values at various EPPS concentrations.

To further investigate the disaggregation efficiency of EPPS across the same temperature range, we performed parallel studies using ThT fluorescence assays. As a control, we first assessed the intrinsic thermal stability of mature amyloid fibrils in the absence of EPPS by monitoring ThT fluorescence intensity over 24 hours at 35°C (Supplementary Figure S9). The ThT intensity remained constant throughout this period, indicating no detectable loss of ThT-binding *β*-sheet signal under these thermal conditions. Subsequently, we evaluated the disaggregation capacity of EPPS by incubating amyloid fibrils with varying concentrations of EPPS (10, 20, 30, and 40 *µ*M) and monitoring ThT fluorescence at 30-minute intervals over 24 hours at each temperature.

Across the temperature range of 30°C to 45°C, we observed a consistent, monotonic decrease in ThT fluorescence intensity over 24 hours for all EPPS concentrations, consistent with a sustained disaggregation process. The reaggregation behavior previously seen at 25°C for 30 *µ*M and 40 *µ*M EPPS, conditions where EPPS predominantly exists in a clustered, supramolecular state, was strongly suppressed at these higher temperatures. The full kinetic spectra for these experiments are provided in Supplementary Figure S10. This transition suggests that the self-associated state of EPPS plays an important role in influencing the amyloid disaggregation pathway. By thermally promoting dissociation of EPPS clusters, the system appears to shift from a regime exhibiting coupled disaggregation–reaggregation behavior toward a regime dominated by net disaggregation (from Regime 2 to Regime 1). Modifying the reaction temperature not only induced a regime transition but also modulated the disaggregation kinetics. We performed a Hill-equation analysis supported by bootstrap analysis ^48,49^ with 500 iterations, to quantify the kinetic parameters *k*_*D*_ and *n*_*H*_. The resulting *k*_*D*_ values exhibited a consistent decrease with increasing temperature across all EPPS concentrations, confirming that the disaggregation process becomes significantly faster at elevated temperatures, as would be expected. Conversely, cooperativity values were constrained within a consistent range across all temperatures, despite exhibiting fluctuations dependent on EPPS concentration. The complete analysis and extracted kinetic parameters for all temperatures are provided in Supplementary Figure S10 (right column). These findings suggest that while temperature enhances the overall rate of disaggregation (*k*_*D*_), the underlying mechanism of the process, as reflected by the cooperativity, remains robust. This is consistent with the possibility that the regime transition is influenced by the availability of EPPS species, rather than a change in the intrinsic amyloid disassembly mechanism.

To characterize the energetics of the overall disaggregation process, we determined apparent activation thermodynamic parameters from the temperature dependence of the measured disaggregation kinetics. Since amyloid disaggregation involves a heterogeneous ensemble of interconverting species and multiple microscopic processes, these parameters describe the effective energetics underlying the overall observed disaggregation kinetics. They should therefore be interpreted as ensemble-averaged descriptors of the observed process, rather than thermodynamic parameters associated with any specific molecular species, conformational transition, or elementary reaction step. We used the disaggregation half-times (*k*_*D*_), derived from the Hill analysis, to construct Arrhenius plots (Figure 5(b)). Since these *k*_*D*_ values represent the half-time of the reaction, we converted them into apparent rate constants (*k*) using *k* = ln(2)*/k*_*D*_ for all thermodynamic calculations. To isolate and characterize the EPPS-induced disaggregation mechanism, we focused our analysis on the disaggregation rate constants across the entire temperature range. By excluding the secondary re-aggregation regime observed only at 25°C, we ensured that the derived thermodynamic parameters primarily reflect the disaggregation pathway. To ensure the statistical robustness of our thermodynamic parameters, we performed a bootstrap resampling analysis ^50^ by perturbing the experimental *k*_*D*_ values within their measured standard deviations (500 iterations), and propagating the uncertainty of our kinetic measurements through the full thermodynamic analysis pipeline. By applying the Arrhenius equation (*E*_*a*_):

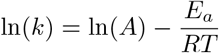

where *k* is the calculated rate constant, *A* is the pre-exponential factor, *R* is the universal gas constant, and *T* is the absolute temperature, we determined the apparent activation energy (*E*_*a*_) across the different EPPS concentrations (Figure 5(d)). The apparent activation energy was extracted from the slope of the linear plot of ln(*k*) versus 1/*T*, where the slope is defined as − *E*_*a*_*/R*. We observed an increase in *E*_*a*_ values with increasing EPPS concentration. Beyond the Arrhenius analysis, we also applied the Eyring–Polanyi equation

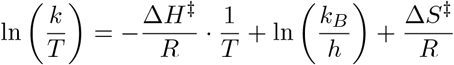

to extract the apparent enthalpy of activation (Δ*H*^*‡*^) and apparent entropy of activation (Δ*S*^*‡*^), providing a thermodynamic description of the transition state ensemble, where *k*_*B*_ is the Boltzmann constant and *h* is Planck’s constant. We generated Eyring plots by plotting ln(*k/T*) versus 1*/T* (Figure 5(c)). The slope of this linear plot provides the apparent enthalpy of activation (Δ*H*^*‡*^) (Supplementary Figure S11(a)), while the intercept allows for the calculation of the apparent entropy of activation (Δ*S*^*‡*^) (Supplementary Figure S11(b)). The Gibbs free energy of activation, Δ*G*^*‡*^, represents the apparent energy barrier associated with the observed ensemble-level process, defined by the relation Δ*G*^*‡*^ = Δ*H*^*‡*^ − *T* Δ*S*^*‡*^. Here, Δ*H*^*‡*^ reflects the apparent energetic contribution associated with the redistribution of aggregate states, while Δ*S*^*‡*^ accounts for the entropic contribution arising from the conformational changes in the ensemble distribution during the observed process.

Notably, the apparent activation enthalpy mirrors the trend of *E*_*a*_, while the apparent activation entropy shows an inverse relationship, i.e., it is more negative at low EPPS concentrations and becomes less negative (shifting toward zero) at higher concentrations. This reciprocal behavior indicates a redistribution between enthalpic and entropic contributions to the activation barrier as the EPPS concentration increases. To investigate this interdependency, we generated apparent enthalpy-entropy compensation plots (Δ*H*^*‡*^ vs. *T* Δ*S*^*‡*^) (Supplementary Figure S11(c)). We observed a linear correlation with a slope of approximately 1, demonstrating strong enthalpy-entropy compensation to the apparent activation barrier. Similar behavior has been reported in other protein aggregation and ligand-binding systems, where enthalpy-entropy compensation acts as a fundamental constraint on the thermodynamic transition state ^51^. ^52^ Furthermore, plotting Δ*H*^*‡*^ against Δ*S*^*‡*^ allowed us to extract the apparent compensation temperature (*T*_*c*_ = 300.15 ± 0.55 K) for the ensemble level process (Figure 5(e)). A *T*_*c*_ value close to room temperature suggests that the amyloid-EPPS system operates near a thermodynamic iso-kinetic-like regime, where changes in enthalpy and entropy partially compensate in determining Δ*G*^*‡*^.

The apparent activation Gibbs free energy (Δ*G*^*‡*^) obtained from the temperature-dependent analysis is ≈ 90 kJ/mol (≈ 21.5 kcal/mol) and remains invariant across the tested EPPS concentration range (Figure 5(f)). This value lies within the range reported for intrinsic A*β*_42_ aggregation and conformational transition barriers in modulator-free systems, ^45^ suggesting preservation of the rate-limiting energetic barrier within the limits of the model. Unlike previous studies typically conducted at millimolar concentrations, ^37,53^ we observe disaggregation and monomer stabilization at micromolar EPPS levels, a regime closer to physiological relevance. Under these conditions, the system behavior is consistent with modulation through redistribution of soluble species. ThT fluorescence control experiments (Supplementary Figure S12) further indicate that EPPS stabilizes monomeric *Aβ*_42_ and inhibits aggregation, consistent with our proposed disaggregation model where the equilibrium is dictated by monomeric interactions (Figure 1). This stabilization may then drive the observed perturbation of the dynamic network, triggering either fragmentation or reaggregation pathways in a concentration-dependent manner. Following disaggregation, the resulting amyloid end-products are long-lived and show no detectable increase in fibrillar signal for up to 5 days (Supplementary Figure S13). This stability, combined with the observed modulation of the structural landscape, strongly supports a monomer sequestration mechanism. While surface-mediated interactions cannot be excluded, the observed compensation behavior suggests a robust and internally consistent energetic description of the disaggregation process.

### Interactions of A*β* and EPPS in macromolecularly crowded environments

Redistribution of soluble species such as that discussed above is expected to be sensitive to environmental constraints such as macromolecular crowding. Cellular environments impose crowding effects that can affect diffusion, accessibility, and fibril organization. To probe this, we introduced PEG2000 as a crowding agent that promotes volume exclusion and stabilizes compact fibril networks. ^54^ TEM imaging of PEG-crowded fibrils revealed pronounced morphological remodeling relative to uncrowded controls. Previously straight filaments (Figure 3(e)) became shorter, bent, and interconnected, forming network-like arrangements (Figures 6(a) and 6(b)). Network domains varied, with the largest spanning 3–4 *µ*m, alongside smaller clusters of 300–1000 nm. In several regions, the fibrils looped or formed partial rings, indicating enhanced lateral association and flexibility under steric constraints. PEG-induced crowding thus promotes fibril bending, lateral interactions, and network formation, producing a more interconnected amyloid architecture.

**Figure 6.**
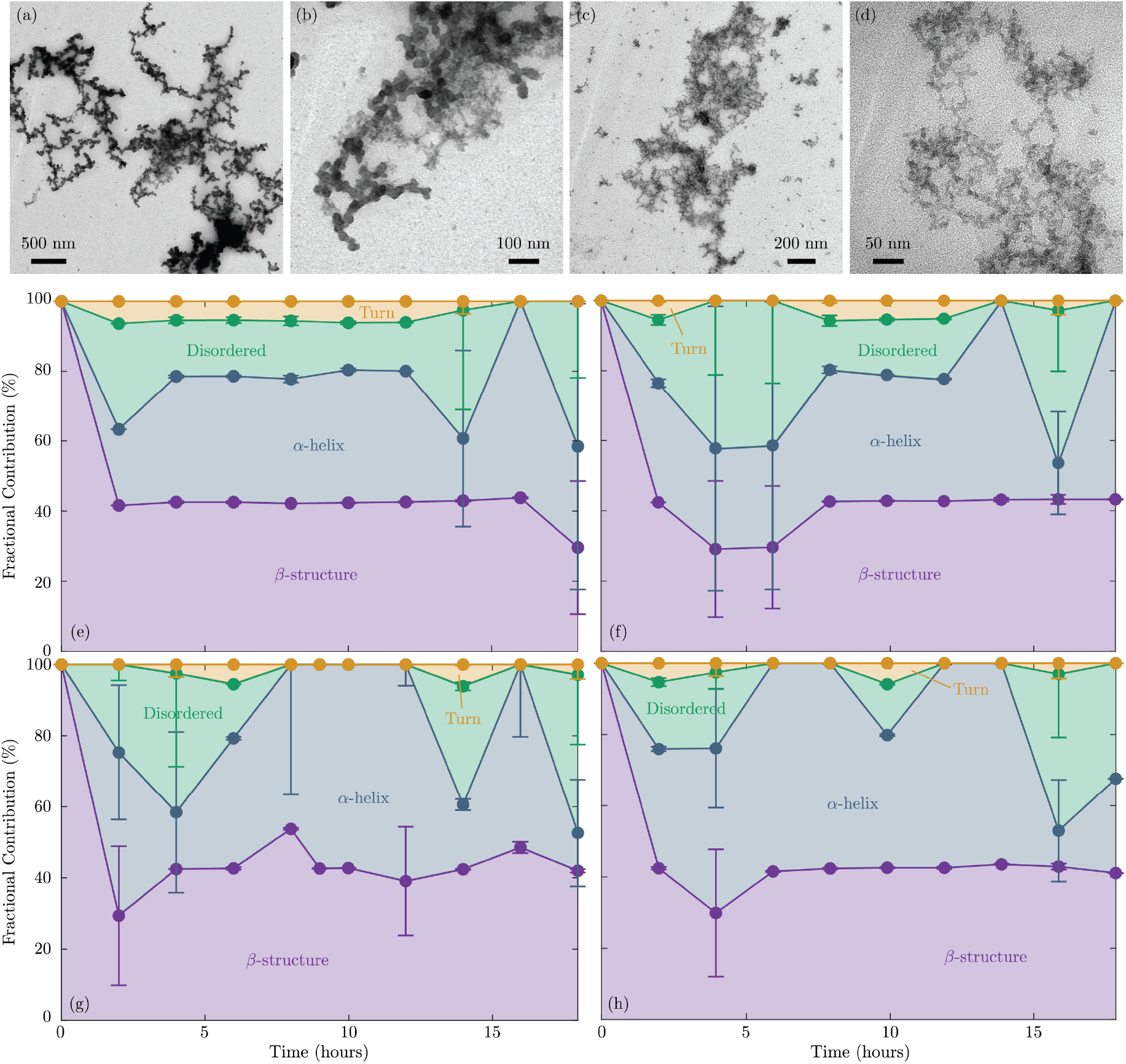
Effect of macromolecular crowding on EPPS induced restructuring of amyloid. (a) and (b) TEM images of PEG-crowded A*β* fibrils. (c) and (d) TEM images of PEG crowded A*β* fibrils after treatment with 20 *µ*M EPPS for 24 h. (e)-(h) FTIR-derived fractional contributions of secondary structural components as a function of time after treatment with EPPS concentrations of (e) 10 *µ*M, (f) 20 *µ*M, (g) 30 *µ*M and (h) 40 *µ*M. Error bars in panels (e)-(h) represent the standard deviation of fractional contributions obtained from three independent replicates.

Treatment with 20 *µ*M EPPS for 24 h largely disrupted large network domains (Figures 6(c) and 6(d)), generating heterogeneous smaller clusters of ~ 200–800 nm while individual fibrils retained elongated morphology. TEM images for treatments with other concentrations of EPPS are shown in Supplementary Figure S14. These observations suggest that EPPS primarily remodels inter-fibrillar contacts, converting large aggregates into smaller assemblies while preserving the intra-fibrillar cross-*β* core. DLS measurements confirmed pronounced size heterogeneity under crowding (Supplementary Figure S15), indicating that although EPPS remodels large clusters, crowding limits complete fibril disassembly.

ThT fluorescence of PEG-crowded fibrils treated with 2–40 *µ*M EPPS exhibited irregular fluctuations similar to PEG-only controls (Supplementary Figure S16), reflecting aggregate-size heterogeneity. The mean ThT signal remained largely unchanged, indicating that individual fibril cross-*β* architecture persisted despite partial network remodeling. These results suggest that EPPS efficacy depends on solvent-exposed *β*-sheet regions, which may be sterically shielded under crowded conditions.

FTIR spectroscopy further provided molecular insight into structural evolution under crowded conditions. The deconvolved Amide I spectra of PEG-treated controls indicated predominantly *β*-sheet-rich fibrillar structure (Supplementary Figure S17(a)), consistent with stabilized amyloid assemblies. Upon EPPS treatment (10-40 *µ*M), a rapid decrease in *β*-sheet content was observed within the first 2 h, reaching ~ 42%, accompanied by transient secondary-structure rearrangements, including *α*-helical signatures and fluctuations in disordered content (Figures 6(e) to 6(h)). A representative deconvolved Amide I FTIR spectrum for EPPS-treated, PEG-crowded A*β* is shown in Supplementary Figure S17(b). Time-resolved measurements at 20 min intervals showed that most of the *β*-sheet reduction occurred within the first 20 min, followed by stabilization of the secondary structure distribution over the remaining observation period (Supplementary Figure S18). No further significant evolution was observed up to 18 h, while fluctuations in the disordered fraction likely reflect transient intermediates during early structural rearrangement.

These observations suggest a heterogeneous response of amyloid assemblies under PEG-induced crowding, where a subset of fibrillar structure undergoes rapid structural reorganization while the remaining cross-*β* core remains comparatively resistant over the experimental timescale. Consistent with ThT measurements, the fibrillar core signal is largely retained, indicating that structural perturbation is primarily associated with accessible or exposed regions rather than complete core disruption. Complementary FTIR measurements performed under PEG-crowding conditions and after extensive washing (Supplementary Figures S19) show that the characteristic PEG-associated spectral features (~1100 cm^−1^) are present during crowding but are not detectable after washing, consistent with removal of PEG from the sample under the conditions examined. These observations are consistent with previous reports describing predominantly excluded-volume interactions of PEG with protein systems. ^55^

Complementary ThT, FTIR, and TEM analyses together provide a coherent picture of EPPS activity under PEG-induced crowding. Although ThT fluorescence remains largely unchanged, FTIR reveals a reduction in *β*-sheet content accompanied by transient redistribution of *α*-helical and disordered structural components, while TEM demonstrates the conversion of extended amyloid networks into smaller, partially reorganized clusters. Collectively, these findings indicate that PEG-induced crowding preserves the fibrillar cross-*β* core while permitting structural remodeling of more accessible regions of the amyloid assemblies over the experimental timescale.

## Conclusion

Our results demonstrate that EPPS modulates A*β*_42_ pathways by promoting the disaggregation of mature fibrils and redistributing protein populations. The invariance of the apparent activation parameters across the tested EPPS concentrations suggests that these population-level shifts occur without measurably altering the effective activation energetics captured by this ensemble-averaged approach. Instead, our data points to a mechanism where EPPS likely acts by destabilizing fibrillar structures and sequestering lower-order metastable species. We found that this remodeling behavior is highly nuanced, sensitive to factors like concentration-dependent small molecule self-association, molecular crowding, and temperature. This is a critical observation because it emphasizes that the efficacy of such modulators is deeply tied to the crowded, dynamic conditions of the actual cellular environment. These findings shift how we might think about therapeutic intervention. Rather than searching exclusively for molecules that act as kinetic blockers to prevent formation, our work suggests that we can achieve meaningful results by targeting the population dynamics of heterogeneous assemblies to reverse existing aggregate states. While the exact microscopic details of how EPPS interacts with A*β*_42_ remain to be fully mapped, this study establishes a foundation for exploring these population-based strategies. Moving forward, the next step will be to bridge these biophysical insights with structural studies to confirm exactly how these remodeled states are formed and maintained in vivo.

## Materials and Methods

### Amyloid fibril preparation

Amyloid beta 42 (A*β*) was purchased from Abcam. Lyophilized amyloid powder (1 mg) was stored at −80°C. To prepare amyloid fibrils, peptide aliquot was dissolved in 10mM NaOH and immediately diluted with phosphate buffered saline (PBS, pH 7.2) to reach 20 *µ*M A*β* solution. The resulting solution was incubated at 37°C with gentle agitation for 24-36 hours to allow the formation of fibrils. Fibril formation was further confirmed using Thioflavin T fluorescence assays and TEM imaging.

### Preparation of EPPS-Amyloid assay

EPPS solutions were prepared by dissolving EPPS in deionized water (pH 6.5) and immediately used. Fully grown amyloid fibrils were mixed with EPPS solution to obtain final EPPS concentrations ranging from 2 *µ*M to 40 *µ*M (step size 2 *µ*M). The final mixtures of EPPS and amyloid had a pH of around 6.8. These mixtures were then subsequently used for spectroscopic measurements and imaging analyses.

### PEG crowded amyloid assay

Cell-mimicking crowded assays were prepared by adding the crowding molecule PEG-2000 (50% w/v) to a 20 *µ*M A*β* fibril solution. PEG:amyloid ratios were varied from 1:10 to 1:2. The mixtures were incubated for 24 hours prior to measurements. The resulting samples were used for spectroscopic measurements and imaging analyses to monitor size changes in amyloid aggregates.

### Fluorescence measurements

Amyloid aggregation was monitored using the Thioflavin T (ThT) fluorescence assay. For the assay, a 20 *µ*M A*β* solution was mixed with 10 *µ*M ThT solution. Three independent samples were prepared, and 50 *µ*L of each was loaded into a microwell plate for fluorescence measurements. Measurements were carried out using a SpectraMax iD5 multimode microwell plate reader (Molecular Devices) with a spectral resolution of 1 nm. The excitation wavelength was 440 nm, and emission was monitored at 485 nm. The experiment was conducted continuously for 24 hours and the readings were taken at 30 minute intervals. Before each reading, the plate was shaken gently for 5 seconds to ensure uniform mixing.

### DLS studies

DLS measurements were performed using Zetasizer advanced ultra (Malvern Panalytical) at 25°C in low volume disposable plastic cuvettes with a 500 *µ*l sample volume. The instrument provides particle resolution of approximately 2:1, capable of distinguising particle populations differing by two fold size. DLS was used to monitor size changes during amyloid growth, EPPS-induced amyloid disaggregation and reaggregation as well as EPPS self-clustering. Measurements were collected over 25 cycles, and the samples were measured 3 to 4 times and averaged to get the mean size distribution. Three independent replicates were analysed for all samples following the same procedure.

### FTIR Measurements

FTIR measurements were performed using a Shimadzu IRTracer-100 to obtain structural information on amyloid intermediates and secondary structures formed during EPPS interactions in both buffer and crowded environments. Experiments were conducted at a spectral resolution of 2 cm^−1^ with 256-scan averaging. Background spectra of the corresponding buffer were subtracted prior to each measurement to ensure accurate analysis. Samples (~2 *µ*L) were measured in attenuated total reflectance (ATR) mode by directly applying the solution onto the diamond crystal at room temperature. Three independent replicates were performed for each condition.

### Zeta potential studies

Zeta potential measurements were carried out using a Zetasizer Advanced Ultra (Malvern Panalytical) at 25°C in low-volume disposable cuvettes with an applied voltage across the electrodes. Samples were prepared in phosphate-buffered saline (PBS, pH 7.4) and equilibrated for 30 s before measurement. For each sample, three measurements were taken and averaged, and three independent replicates were performed. Measurements were used to monitor changes in the surface charge of amyloid fibrils during EPPS-induced disaggregation and reaggregation.

### Transmission Electron Microscopy

For negative stain TEM, 5 *µ*L of each sample was incubated for 60 s on a freshly glow discharged copper grid covered with a continuous carbon film. The grids were blotted with a filter paper and stained by applying a drop of 1% uranyl acetate on grids for 60 s, blotted with filter paper and air-dried for 10 min before TEM viewing. The grids were imaged using FEI Tecnai 120 keV with a digital CCD Eagle camera 4k × 4k (Thermo Fisher Scientific,USA).

## Author contributions

RVN and MRF designed the study. RVN performed the experiments, analyzed the data, and wrote the manuscript. BNT performed TEM structural characterization. ANP contributed to data interpretation and discussion. MRF supervised the research, refined the data visualization, and contributed to manuscript revision. All authors reviewed and approved the final manuscript.

## Acknowledgments

This research is supported by the Ministry of Education, Singapore, under its Research Centre of Excellence award to the Institute for Digital Molecular Analytics & Science, NTU (IDMxS, grant: EDUNC-33-18-279-V12). The authors acknowledge the use of NISB Cryogenic Electron Microscopy Platform at NTU Institute of Structural Biology (NISB), Nanyang Technological University.

