## Supplementary Material for "Small-molecule-induced species redistribution drives amyloid remodeling along a conserved kinetic landscape"

### 1 Amyloid beta growth - ThT fluorescence assay and DLS measurements

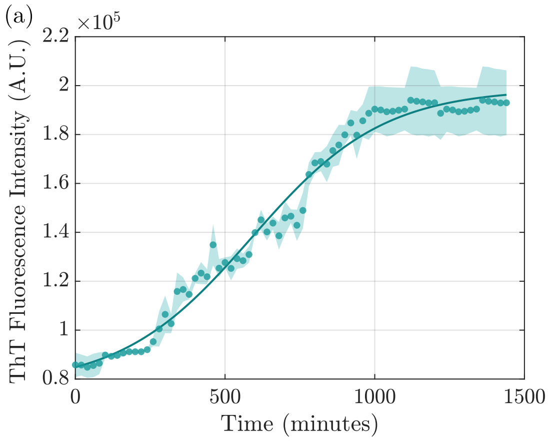


Figure S1: (a) ThT fluorescence versus incubation time showing amyloid beta growth kinetics. The apparent rate constant ($k$), lag time ($t_{\mathrm{lag}}$), and half-time ($t_{1/2}$) are extracted from the sigmoidal fit as 0.0046 min$^{-1}$, 587 min, and 150 min, respectively. (b) Dynamic light scattering (DLS) data showing the change in particle size during incubation, reflecting aggregation behavior over time. The solid line in (a) indicates the sigmoidal fit, and the shaded bands in both panels represent the standard deviation from three independent replicates.

### 2 Steady state ThT data during EPPS treatment

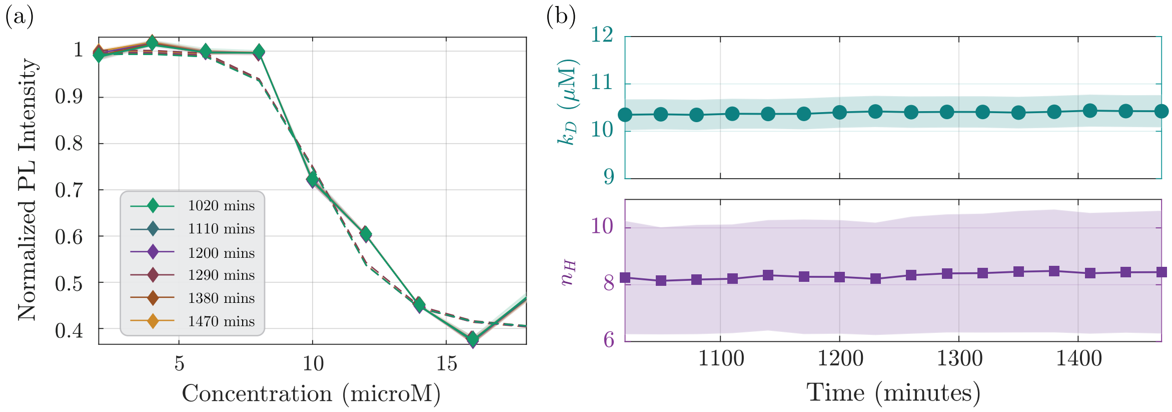


Figure S2: (a) ThT intensity as a function of EPPS concentration at different incubation times during the steady phase (900-1500 minutes, shown as solid curves of different colors). (b) Time-dependent plots of $k_D$ and $n_H$ during the steady state obtained from sigmoidal fit (dashed line). The shaded band in (a) and (b) indicates the standard deviation obtained from 3 independent replicates.

### 3 Binding isotherm

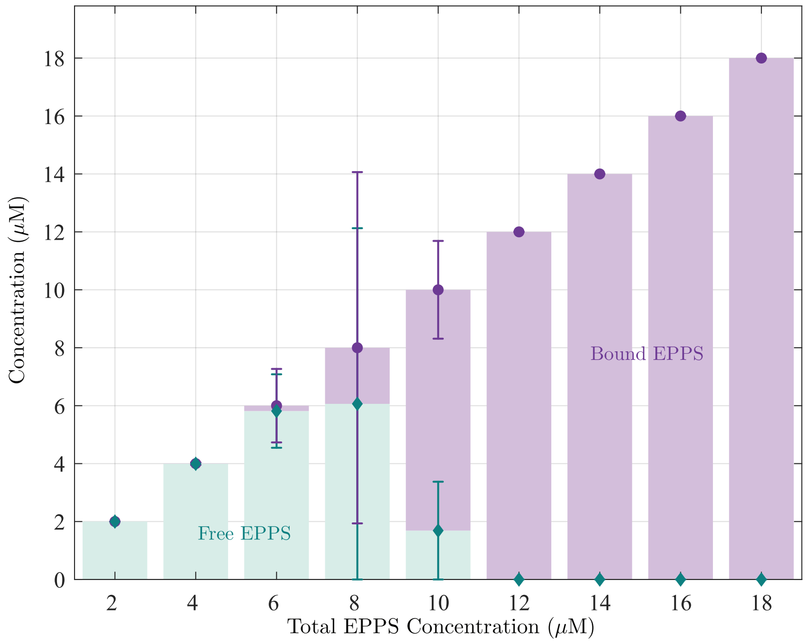


Figure S3: Free and bound EPPS concentrations as a function of total EPPS concentration, derived from Hill fitting of ThT steady-state kinetic data. Error bars represent the standard deviation of calculated free and bound EPPS concentrations from three independent replicates.

### 4 FTIR measurements

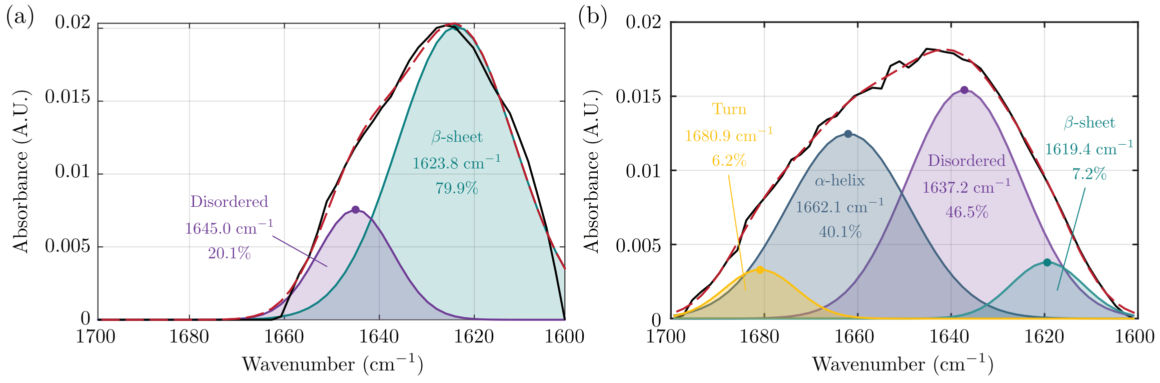


Figure S4: (a) FTIR spectra of amide I region of fully grown amyloid beta fibrils. Deconvolution using pseudo-Voigt fit shows $\beta$-sheet as the major component with peak area of ∼80% along with the disordered peak with peak area of ∼20%. (b) FTIR spectra of the amide I region of A$\beta$ after 4 hours of treatment with 10 $\mu$M EPPS. Deconvolution using a pseudo-Voigt fit reveals a marked reduction in $\beta$-sheet content, accompanied by the emergence of new secondary structural components, consistent with EPPS-induced disaggregation of A$\beta$ fibrils.

### 5 TEM – EPPS treated A$\beta$ – multi-day incubation

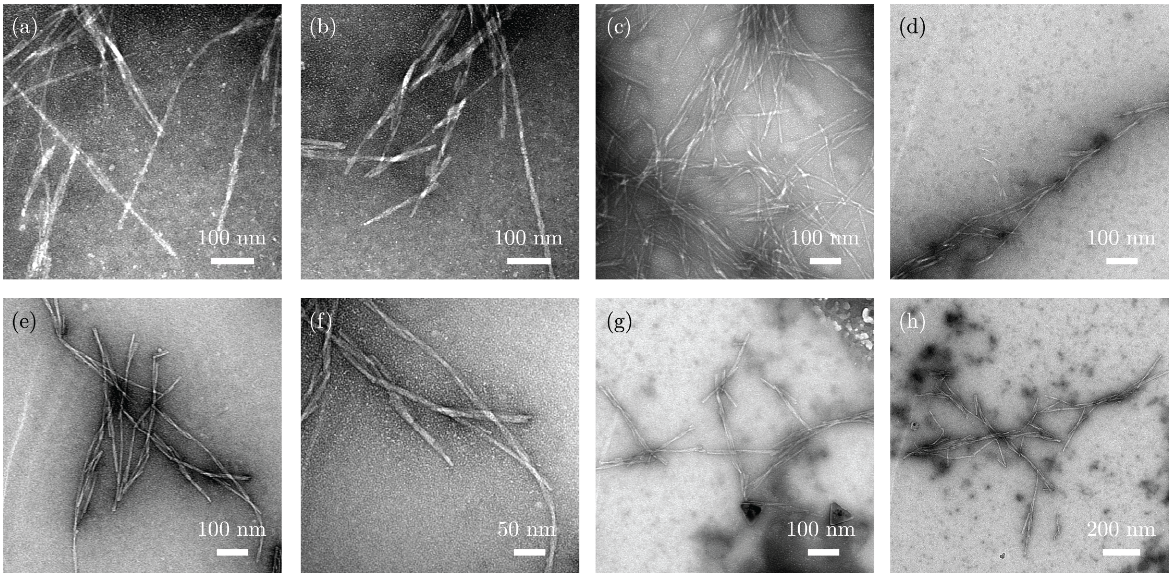


Figure S5: Negative-stain TEM images of A$\beta_{42}$ fibrils under different EPPS concentrations. (a) and (b) 30 $\mu$M EPPS treated sample incubated for 4 days. (c) and (d) 30 $\mu$M EPPS treated sample incubated for 6 days. (e) and (f) 40 $\mu$M EPPS treated sample incubated for 4 days. (g) and (h) 40 $\mu$M EPPS treated sample incubated for 6 days

### 6 Estimation of EPPS cluster fraction from DLS measurements

To estimate the percentage of free monomers and clustered species from DLS intensity data, we applied scattering theory principles to convert intensity distributions into approximate number distributions. For clusters larger than 60 nm, we used Mie theory, which accounts for complex size-dependent scattering effects. According to Mie theory, the scattering efficiency $Q_{\text{sca}}$ is calculated using the scattering coefficients $a_n$ and $b_n$ as follows [1]

$$Q_{\text{sca}} = \frac{2}{x^2} \sum_{n=1}^{\infty} (2n+1) \left( |a_n|^2 + |b_n|^2 \right),$$

where the size parameter $x = \frac{2\pi r}{\lambda}$, $r$ is the particle radius and $\lambda$ is the wavelength in the host medium. The coefficients $a_n$ and $b_n$ are given by:

$$a_n = \frac{m\psi_n(mx)\psi_n'(x) - \psi_n(x)\psi_n'(mx)}{m\psi_n(mx)\xi_n'(x) - \xi_n(x)\psi_n'(mx)},$$

$$b_n = \frac{\psi_n(mx)\psi_n'(x) - m\psi_n(x)\psi_n'(mx)}{\psi_n(mx)\xi_n'(x) - m\xi_n(x)\psi_n'(mx)},$$

where $m = \frac{n_{\text{particle}}}{n_{\text{medium}}}$ is the relative refractive index, $\psi_n$ and $\xi_n$ are the Riccati-Bessel functions, and primes denote derivatives with respect to the argument. These coefficients are calculated numerically. The scattered intensity is proportional to the product of the number of particles $N$, the scattering efficiency $Q_{\text{sca}}(d)$, and the geometric cross-section ($\propto d^2$), expressed as

$$I(d) \propto N Q_{\text{sca}}(d) d^2$$

where $d = 2r$ is the particle diameter Rearranging this relation allowed us to estimate the raw number distribution from the DLS data as:

$$N_{\text{DLS}}(d) \propto \frac{I(d)}{Q_{\text{sca}}(d) d^2}.$$

To convert this to an absolute number distribution, we calibrated the data using DLS measurements of 500 nm latex beads of known concentration, where the total number $N_{\text{latex}}$ was used to calculate a scaling factor:

$$\text{scaling factor} = \frac{N_{\text{latex}}}{\sum_d N_{\text{DLS}}(d)}.$$

The final calibrated number distribution for clusters larger than 60 nm is then:

$$N(d) = \text{scaling factor} N_{\text{DLS}}(d).$$

For clusters smaller than 60 nm, we assumed the Rayleigh scattering regime ($d \ll \lambda$), where intensity per particle follows the [1]:

$$I_{\text{monomer}} \propto d^6,$$

which can be written as $I_{\text{monomer}} = C d^6$ where $C$ is a proportionality constant. Using DLS data from a 1 nM concentration solution of EPPS as the reference monomer data, we calculated the scaling factor for the smaller clusters. The known monomer concentration is:

$$N_{\text{ref}} = c_{\text{ref}} V_{\text{ref}} N_{\text{A}},$$

where $N_{\text{A}}$ is Avogadro's number. Using the peak intensity $I_{\text{peak}}$ from the reference DLS data, we obtained the scaling factor $C$ by:

$$C = \frac{I_{\text{peak}}}{N_{\text{ref}} d^6}.$$

This allowed us to calculate the expected number of monomers $N_{\text{exp}}$ in the sample as:

$$N_{\text{exp}} = \frac{I_{\text{peak}}}{C d^6}.$$

Through this combined approach, using Mie theory for larger clusters calibrated with latex beads and Rayleigh approximation for smaller clusters calibrated with monomer EPPS data, we translated intensity data into an approximate number distribution of free monomers versus clustered species. A monomer hydrodynamic diameter of 1 nm was assumed; all particles with apparent sizes greater than 1 nm were considered as monomers incorporated within clusters. This provided insights into the fraction of EPPS molecules in monomeric and aggregated states as a function of concentration.

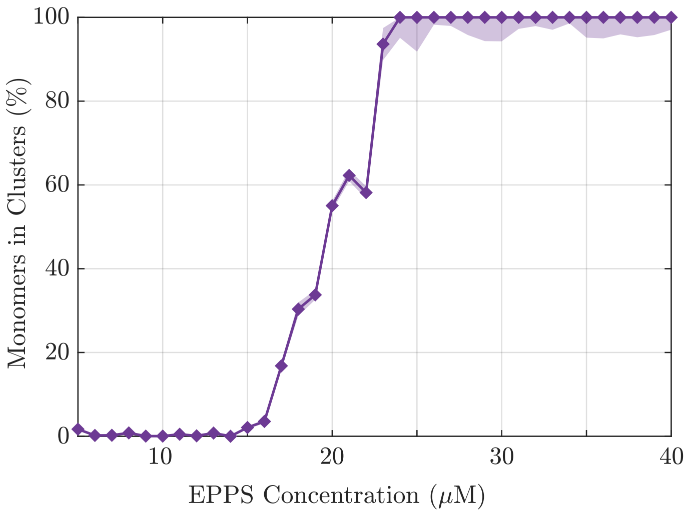


Figure S6: Percentage of EPPS incorporated into clusters as a function of EPPS concentration, extracted from DLS intensity distributions after conversion to number distributions using Rayleigh - Mie scattering models. The fraction of EPPS incorporated into clusters remains nearly zero below ~15 $\mu$M, beyond which it rises sharply, reaching close to 100% around ~25 $\mu$M. This concentration-dependent transition aligns closely with the regimes defined earlier.

### 7 Effect of ionic strength on EPPS dynamic clustering

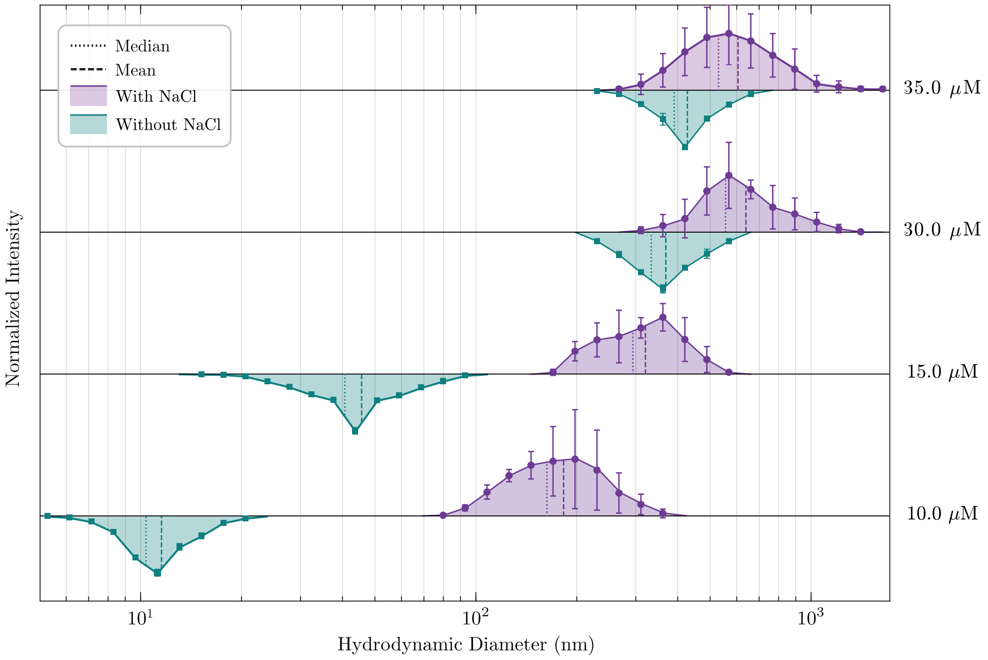


Figure S7: DLS analysis of EPPS (10-35 $\mu$M) dynamic clustering under varying ionic strength via 200 mM NaCl addition. Comparison between the EPPS size distribution shows an increase in the average cluster size formed upon addition of NaCl. Error bars represent standard deviation from three independent measurements. Dashed lines and dotted lines represent the mean values and median values, respectively.

### 8 Effect of HEPES treatment on amyloid beta disaggregation

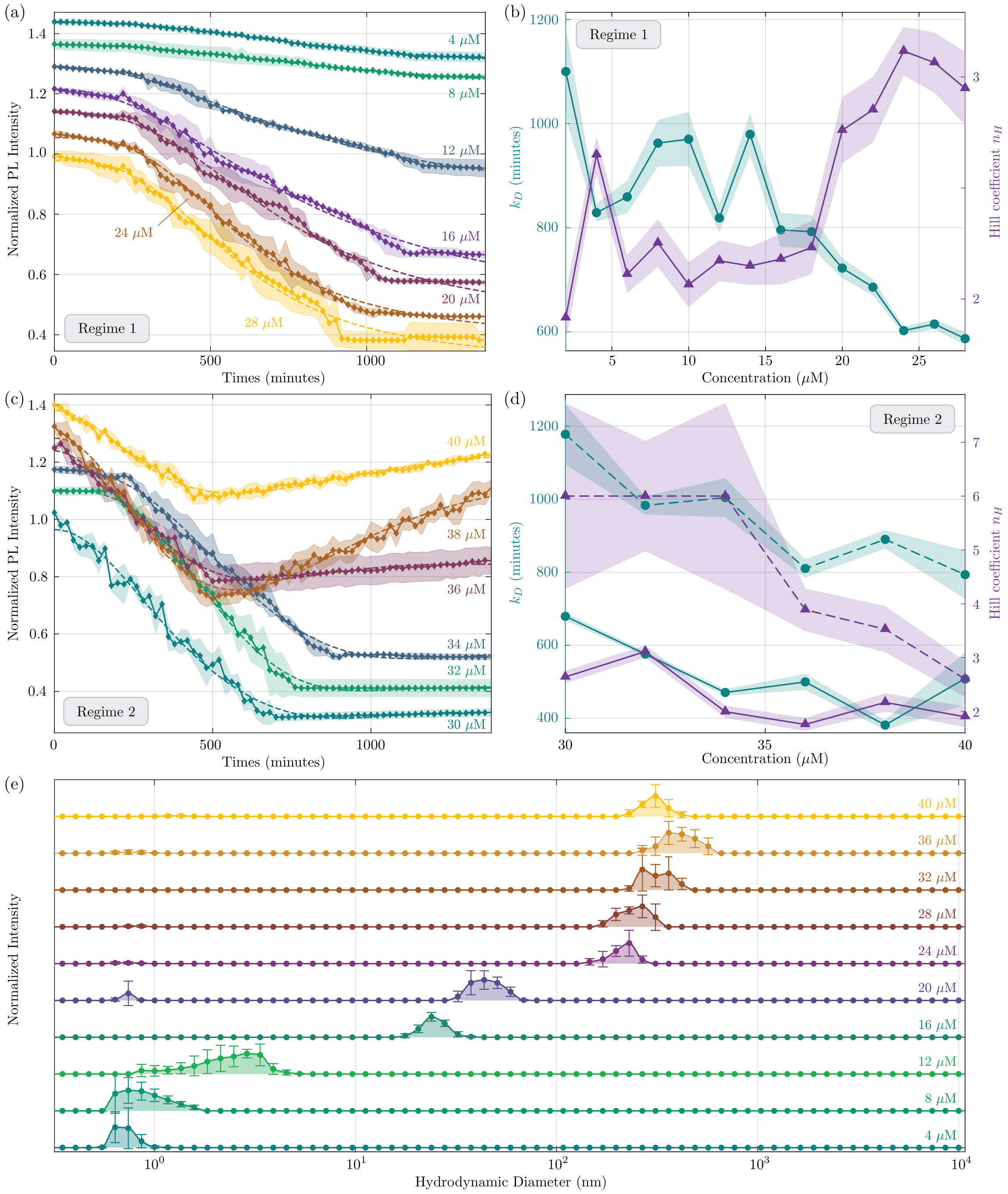


Figure S8: HEPES treatment effects on A$\beta$ aggregation and clustering. (a) ThT PL intensity profiles for Regime 1, (b) ThT PL intensity profiles for Regime 2, (c) Regime 1: $k_D$ and $n_H$ values extracted from Hill fit, (d) Regime 2: $k_D$ and $n_H$ values extracted from double Hill fit, (e) DLS plots showing dynamic clustering of HEPES at various concentrations.

### 9 Temperature-dependent size evolution of EPPS clusters

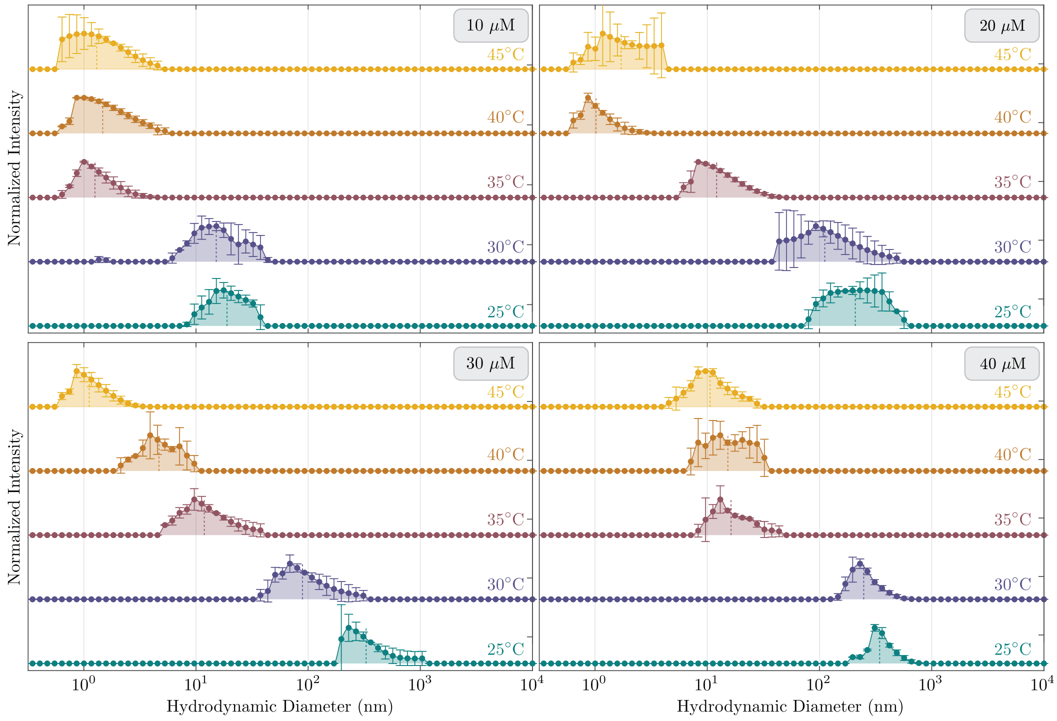


Figure S9: Full DLS spectra of EPPS clusters at various temperatures (25°C to 45°C) for 10 $\mu$M, 20 $\mu$M, 30 $\mu$M, and 40 $\mu$M EPPS concentrations. Data are presented as mean values, with error bars representing the standard deviation from three independent replicate measurements.

### 10 Stability of amyloid fibrils at 35°C

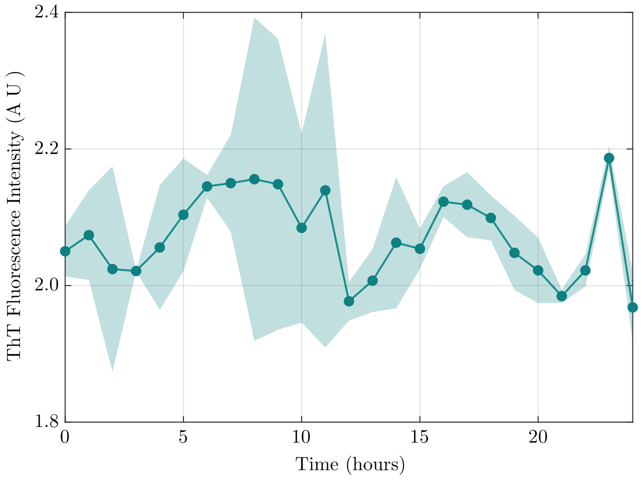


Figure S10: Control experiment assessing the stability of amyloid fibrils using ThT fluorescence at 35°C. Emission intensity was monitored over 24 hours, during which no significant change was observed, indicating that the fibrils remain stable under these conditions.

### 11 Temperature-dependent EPPS-induced A$\beta$ disaggregation kinetics

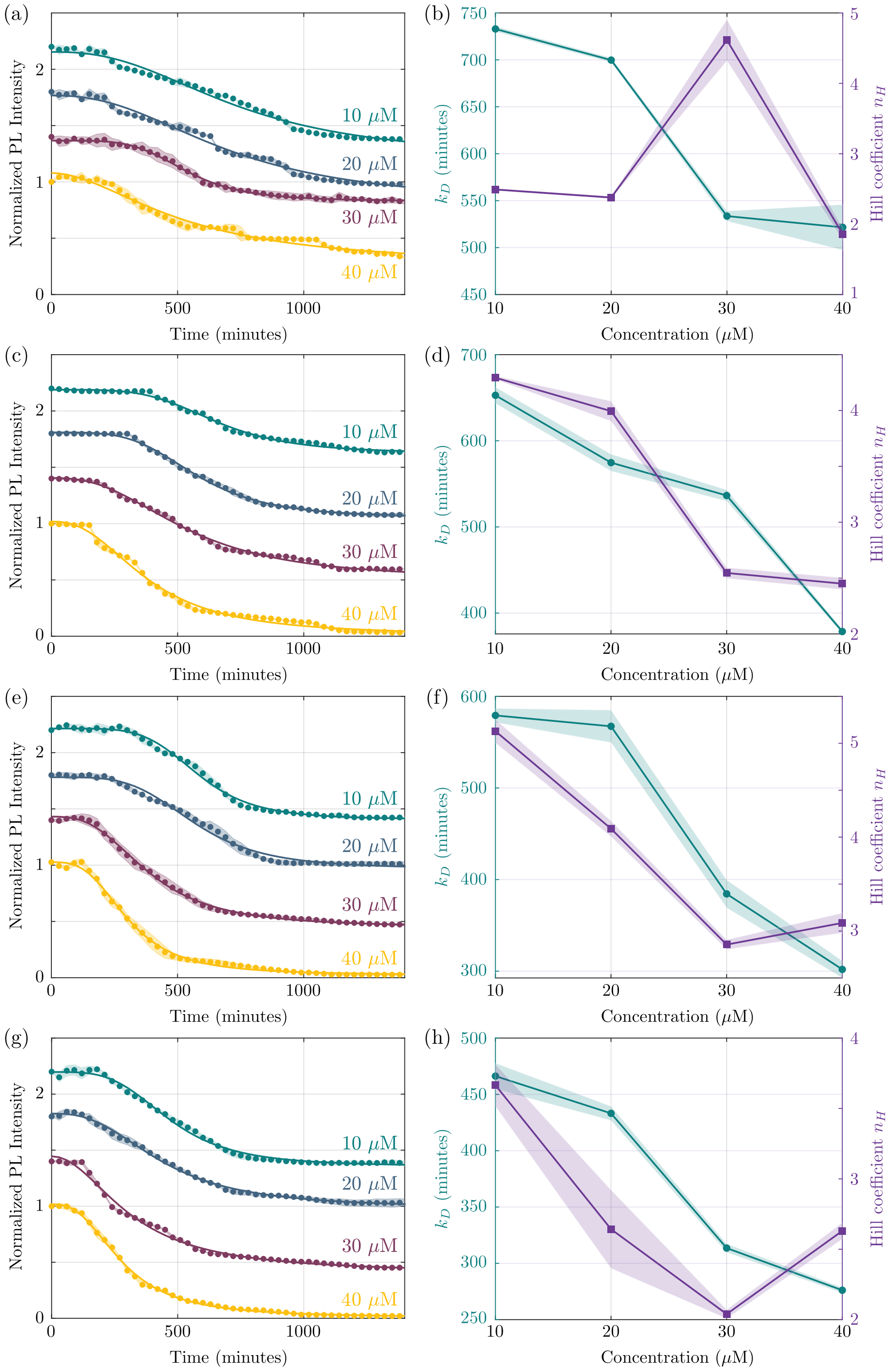


Figure S11: Time-dependent ThT fluorescence intensity for amyloid samples treated with EPPS at four different concentrations ((a) 10, (c) 20, (e) 30, and (g) 40 $\mu$M), measured at 30°C, 35°C, 40°C, and 45°C, respectively with corresponding Hill fit parameters $k_d$ and $n_H$ ((b), (d), (f) and (h)).

### 12 Thermodynamic analysis of EPPS-induced A$\beta$ disaggregation

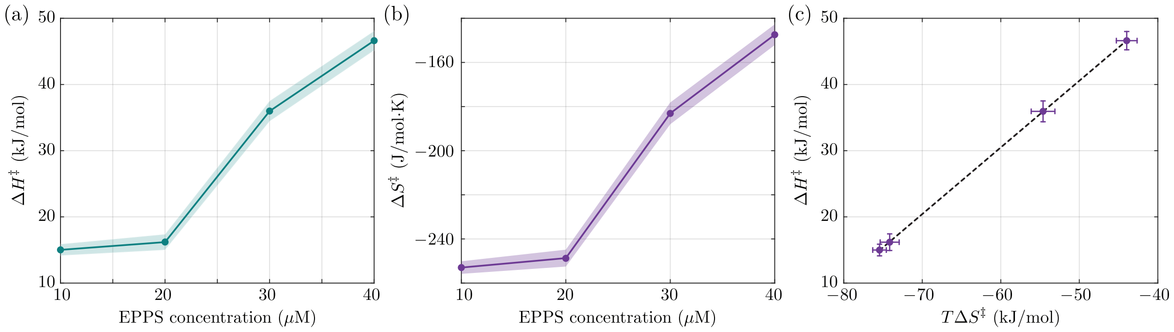


Figure S12: Extracted thermodynamic parameters from Eyring-Polanyi plots. (a) Apparent activation enthalpy ($\Delta H^{\ddagger}$) as a function of EPPS concentration. (b) Apparent activation entropy ($\Delta S^{\ddagger}$) as a function of EPPS concentration. (c) Correlation between $\Delta H^{\ddagger}$ and $T\Delta S^{\ddagger}$, yielding a slope of 1.01 and demonstrating enthalpy-entropy compensation.

### 13 Amyloid monomer stabilization by EPPS

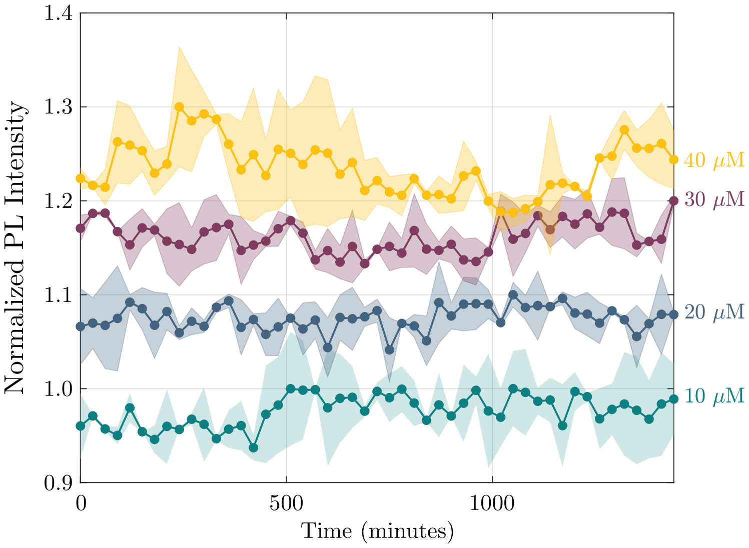


Figure S13: Time-dependent ThT fluorescence intensity of amyloid monomers in the presence of EPPS at concentrations of 10, 20, 30, and 40 $\mu$M. ThT fluorescence was monitored to assess potential amyloid aggregation over time. No significant increase in fluorescence intensity was observed at any EPPS concentration, indicating the absence of detectable fibril formation. These results suggest that EPPS stabilizes the monomeric state and suppresses amyloid aggregation under the experimental conditions.

#### 14 5-day ThT fluorescence study of EPPS-treated A$\beta$

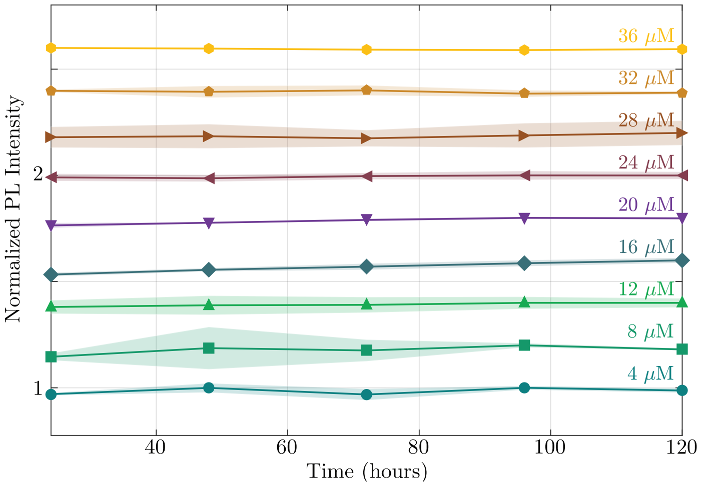


Figure S14: Stacked normalized ThT fluorescence intensity as a function of time for different EPPS concentrations over 5 days. Each trace is vertically offset for clarity and normalized to its initial maximum intensity. The data show no significant increase in ThT signal over time, indicating the absence of detectable secondary aggregation or re-nucleation under the tested conditions. Shaded regions represent variability across 3 replicate measurements.

#### 15 TEM images of EPPS-treated PEG crowded A$\beta$

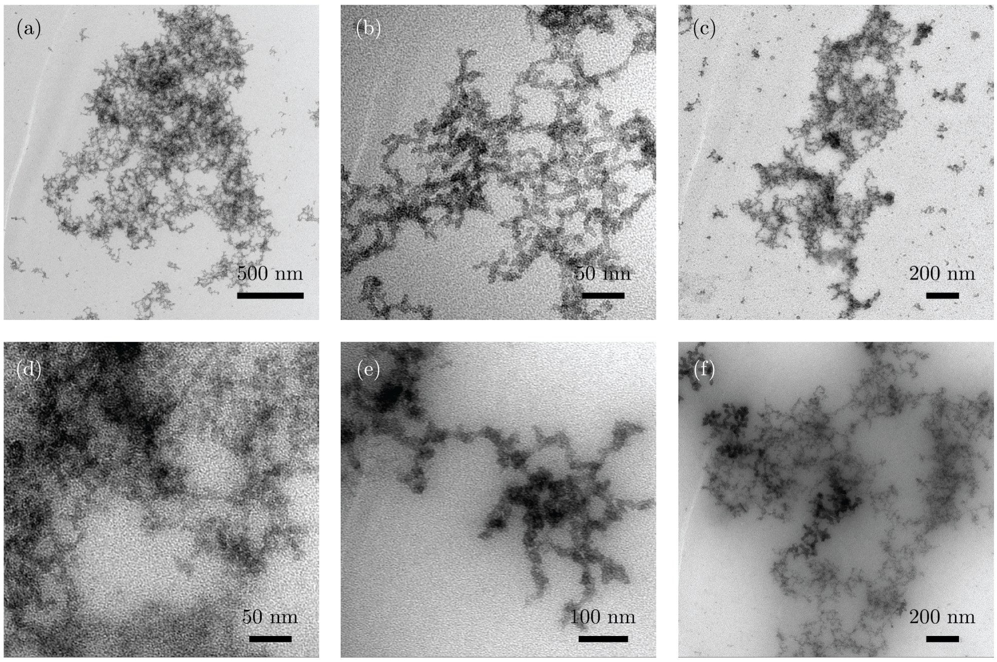


Figure S15: Negative-stain TEM images of PEG crowded A$\beta_{42}$ fibrils treated with various concentrations of EPPS for 24 hours. (a) and (b) 10 $\mu$M EPPS treated sample. (c) and (d) 20 $\mu$M EPPS treated sample. (e) and (f) 30 $\mu$M EPPS treated sample.

#### 16 DLS analysis of amyloid crowding induced by PEG2000

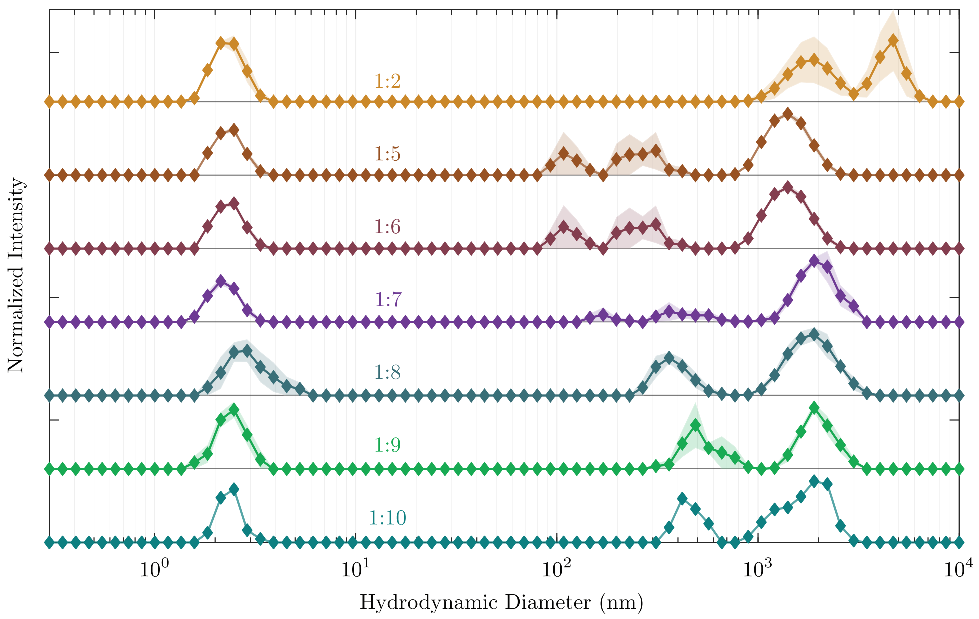


Figure S16: Dynamic light scattering (DLS) intensity versus particle size for amyloid $\beta$ (A$\beta$) in the presence of PEG2000 at various crowding ratios. Shaded bands represent the standard deviation from three replicate measurements. The DLS data show large variations in size distribution, as evident from the standard deviation, indicating that PEG2000 crowding results in heterogeneous clusters within the sample.

#### 17 ThT fluorescence of EPPS treated PEG-crowded fibrils

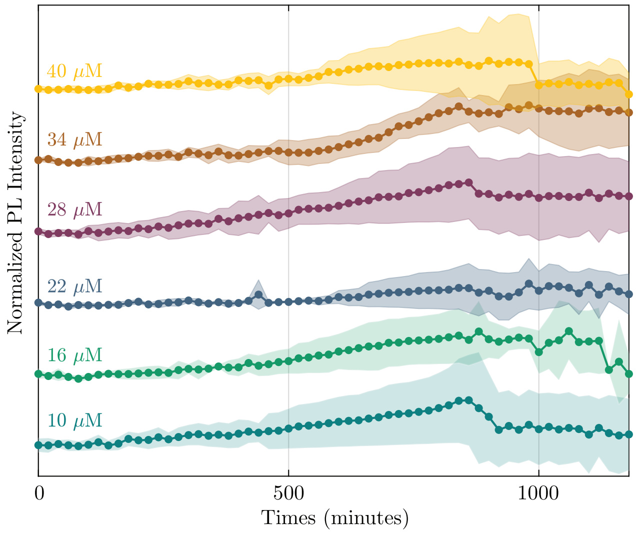


Figure S17: ThT fluorescence intensity as a function of time for PEG-crowded A$\beta$ fibrils treated with varying concentrations of EPPS. Shaded regions indicate the standard deviation of ThT intensity from three independent measurements.

### 18 FTIR spectrum of PEG-crowded A$\beta$

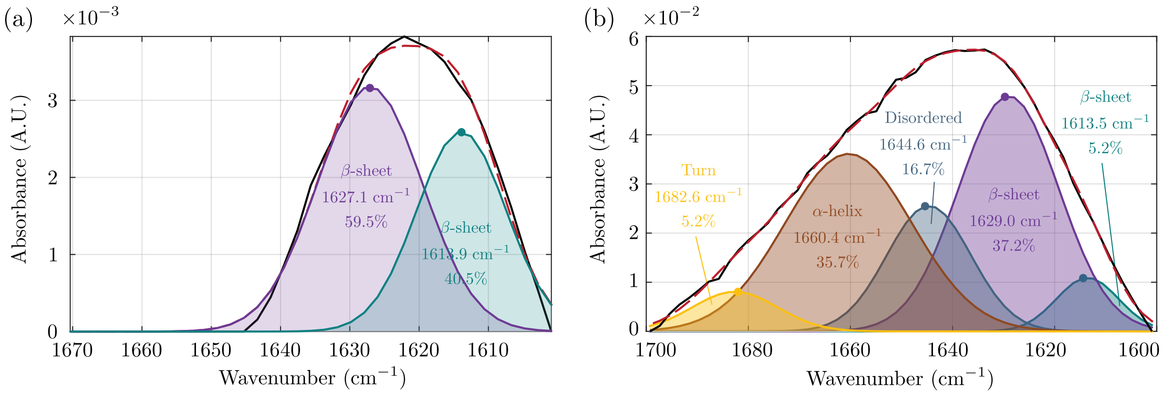


Figure S18: (a) FTIR spectrum of PEG-crowded A$\beta$ fibrils. The amide I region reveals that nearly the entire sample adopts a $\beta$-sheet conformation, indicating a highly ordered fibrillar structure under macromolecular crowding conditions. (b) FTIR spectrum of PEG-crowded A$\beta$ fibrils treated with 10 $\mu$M EPPS for 4 h. The amide I region reveals a substantial decrease in $\beta$-sheet content to approximately 40%, accompanied by the emergence of an $\alpha$-helical peak as the dominant secondary structure, indicating pronounced structural reorganization upon EPPS treatment under crowding.

#### 19 FTIR analysis of PEG crowded EPPS-treated A$\beta$ during early incubation intervals

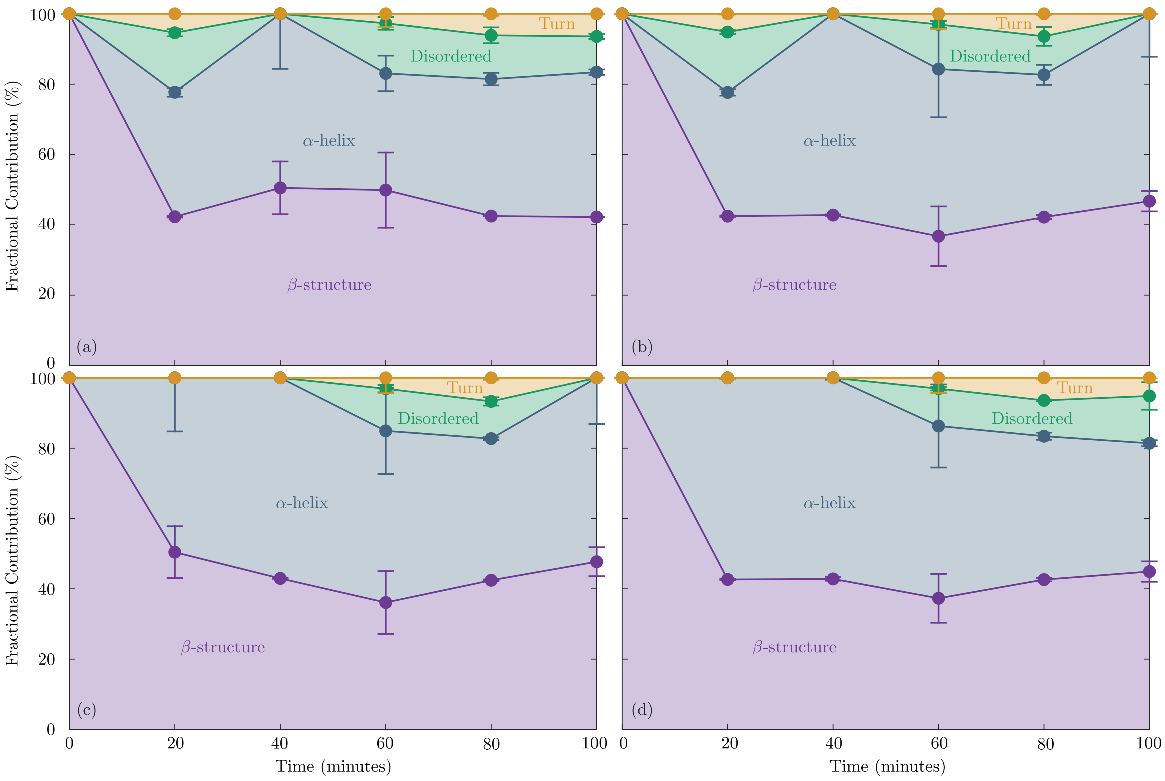


Figure S19: FTIR disaggregation analysis showing concentration-dependent changes in A$\beta$ secondary structures upon treatment with (a) 10 $\mu$M , (b) 20 $\mu$M, (c) 30 $\mu$M and (d) 40 $\mu$M EPPS for 100 minutes. Fractional contribution of $\beta$ and other secondary structures during EPPS treatment are shown as a function of time for the early 100 minute interval.

#### 20 FTIR analysis of PEG – amyloid interaction

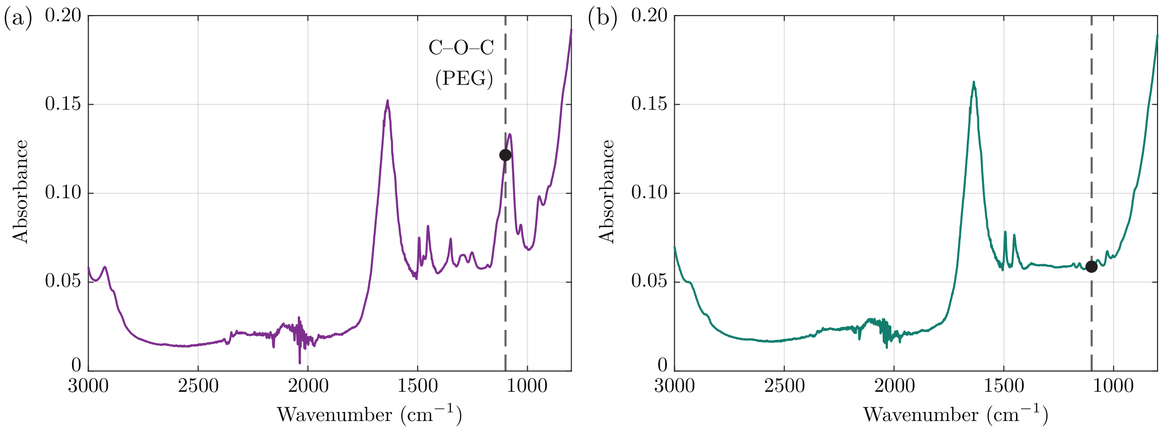


Figure S20: FTIR spectra of PEG-crowded A$\beta$ before and after washing. (a) Spectrum recorded under PEG-crowding conditions showing a characteristic PEG-associated band near $\sim 1100$ cm$^{-1}$ (C–O–C stretching). (b) Spectrum recorded after washing, where PEG-associated spectral features are not detectable, consistent with removal of PEG from the sample under the experimental conditions.
